# Infection dynamics of endosymbionts that manipulate arthropod reproduction

**DOI:** 10.1101/2024.06.23.600294

**Authors:** Franziska A. Brenninger, Roman Zug, Hanna Kokko

## Abstract

A large proportion of arthropod species are infected with endosymbionts, some of which selfishly alter host reproduction. The currently known forms of parasitic reproductive manipulations are male-killing, feminization, cytoplasmic incompatibility, parthenogenesis induction and distortion of sex allocation. While all of these phenomena represent adaptations that enhance parasite spread, they differ in the mechanisms involved and the consequent infection dynamics. We focus here on the latter aspect, summarizing existing theoretical literature on infection dynamics of all known reproductive manipulation types, and completing the remaining knowledge gaps where dynamics have not been modelled yet. Our unified framework includes the minimal model components required to describe the effects of each manipulation. We establish invasion criteria for all potential combinations of manipulative endosymbionts, yielding predictions for an endosymbiont’s increase from rarity within a host population that is initially either uninfected or infected with a different symbiont strain. We consider diplodiploid and haplodiploid hosts, as the mechanisms as well as the infection dynamics of reproductive manipulations can differ between them. Our framework reveals that endosymbionts that *a priori* have the best invasion prospects are not necessarily the most commonly found ones in nature; priority effects play a role too, and cytoplasmic incompatibility excels in this regard. As a whole, considerations of the ease with which a symbiont spreads have to be complemented with knowledge of how easy it is to achieve a particular manipulation, and other factors influencing host switches.

## I. Introduction

Symbiosis is defined as a permanent or prolonged association of different species (de Bary, 1879). In endosymbiosis, the symbiont lives inside the body of its host organism, often within host cells.

Interactions between symbionts and their hosts can switch between parasitism and mutualism even within one species pair (Canestrari *et al*., 2014, Drew *et al*., 2021), as the interests of the two parties are somewhat, but typically not completely, aligned.

A symbiont’s fitness can be increased by manipulating its host’s behaviour: as particularly dramatic examples, *Toxoplasma* increases its transmission from rodents to feline predators by inhibiting the anti-predatory behaviours of the prey (as a side effect, human hosts may increase risk-taking, Flegr *et al*., 2002, Johnson *et al*., 2018), and *Ophiocordyceps* fungi cause ants to choose locations that maximize fungal fitness upon ant death (de Bekker *et al*., 2021). In other cases, the manipulations directly target host reproduction. This is the case for many endosymbionts of arthropods (Hurst & Frost, 2015), the focus of this review.

Endosymbiotic infections of arthropod hosts are very common: Weinert *et al*. (2015) estimate *Wolbachia* to infect 52% of arthropod species, while *Rickettsia* and *Cardinium* reach 24% and 13%, respectively. These bacteria (and additional ones such as *Spiroplasma, Arsenophonus*), as well as some non-bacterial endosymbionts (some microsporidians: Hurst & Frost, 2015; Kageyama *et al*., 2012; and viruses: Kageyama *et al*., 2017*b*; Nagamine *et al*., 2023) manipulate the reproduction of their hosts, which may be insects, arachnids (Curry *et al*., 2015) or crustaceans (Cordaux *et al*., 2001; Moret *et al*., 2001).

There are multiple types of reproductive manipulation (Figure 1), and a given symbiont may induce different types of manipulation depending on host species (Sasaki *et al*., 2002; Jaenike, 2007).

**Figure 1:**
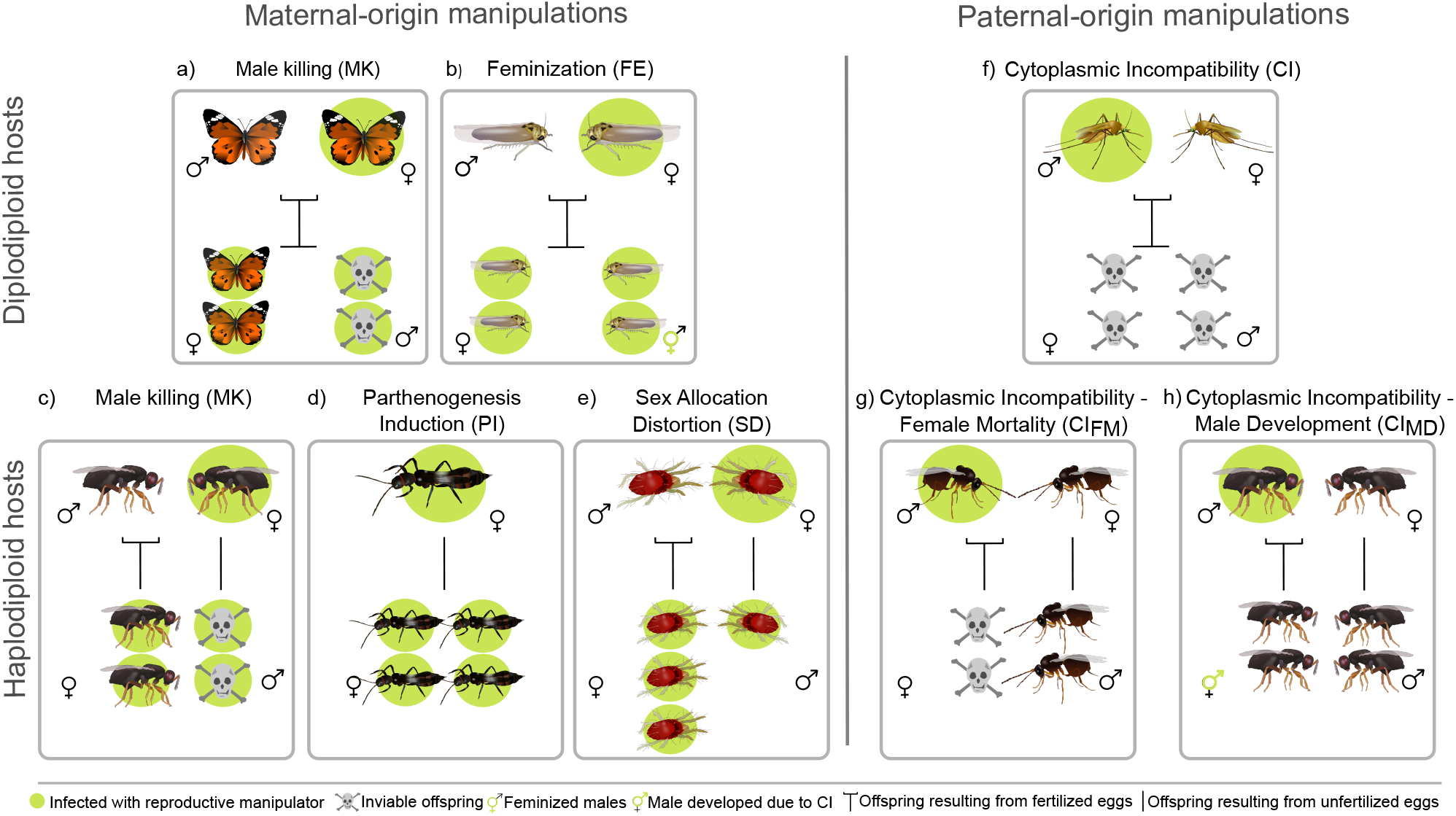
Reproductive manipulations found in arthropod hosts. Schematic representation of known reproductive manipulations: a) & c) male-killing (MK), b) feminization (FE), d) parthenogenesis induction (PI), e) sex allocation distortion (SD) (all these manipulation types originate from maternal infection); f) cytoplasmic incompatibility (CI), g) female mortality CI, and h) male development CI (all CI types originate from paternal infection). Manipulations are further classified into those that are found in diplodiploid host species (top row: a, b, f) and those found in haplodiploid host species (bottom row: c, d, e, g, h). Green circles indicate infection with the respective endosymbiont. Individuals in the upper part of each box form the parental generation and individuals in the lower part represent offspring. If the line connecting parents to their offspring is fork-shaped, the offspring is produced sexually, otherwise it is produced uniparentally. This scheme assumes perfect transmission efficiency *T* and manipulation efficiency *β* except for SD, where *β* < 1, highlighting the result that offspring sex ratio is biased towards females, without necessarily leading to an all-female brood. In comparison, we assume that crosses between uninfected individuals will result in an offspring sex-ratio of 1:1.

Simultaneously, different symbiont strains can induce the same manipulation (Hurst & Frost, 2015). Infection may feature mutualistic and parasitic aspects concurrently (‘Jekyll-and-Hyde symbioses’, e.g., Jiggins & Hurst, 2011; Zug & Hammerstein, 2015). Benefits to the host include the endosymbiont providing additional nutrients or resistance against parasites (Zug & Hammerstein, 2015), while manipulations of host reproduction often feature conflict where the selfish interest of the endosymbiont reduces host fitness (Hurst & Frost, 2015).

The rationale behind reproductive manipulation is the maternal transmission of endosymbionts, where eggs represent the only viable transmission route. A male is thus a dead-end for the endosymbiont, and selection favours manipulations that increase the proportion of infected females in the host population (Engelstädter & Hurst, 2009; Hurst & Frost, 2015). We abbreviate the currently known manipulation types — male-killing, feminization, parthenogenesis, distortion of sex allocation and cytoplasmic incompatibility — as MK, FE, PI, SD and CI, respectively. Some of these behave differently in haplodiploid hosts and/or have subcategories that differ subtly in their dynamics, bringing the total number of distinct cases we consider to 8 (Figure 1).

The molecular mechanisms involved are better understood for some cases (e.g., *Wolbachia*-induced CI, Hochstrasser, 2023) than others, where candidate genes are being discovered (e.g., MK, Perlmutter *et al*., 2019; Kageyama *et al*., 2023; PI, Fricke & Lindsey, 2023; Li *et al*., 2024). Since a recent review of the molecular aspects is available (Benetta *et al*., 2021), we here only briefly outline the general effects of each manipulation and spend the majority of our effort guiding the reader through the theoretical work that has addressed the infection dynamics of each case.

## II. Types of manipulation

Here we give a brief overview of the rules governing different types of reproductive manipulation, without yet commenting on the consequences for infection dynamics. For a more detailed review on manipulation types, we refer the reader to Hurst & Frost (2015).

### (1) Maternal-origin manipulation

In the first major category, maternal-origin manipulation (Figure 1a-e), the manipulation occurs when the mother is infected; the infection status of her mate has no effect. Maternal-origin manipulations create female-biased offspring sex ratios, and in some cases also reduce the total number of viable offspring.

Under **male-killing** (MK, Figure 1a, c), infected male progeny of a female host die during embryonic development (early MK) or in larval and pupal stages (late MK, which we do not explicitly consider here) (Hurst & Frost, 2015). Infected female offspring develop normally. This manipulation can operate in both diplodiploid (Figure 1a) and haplodiploid (Figure 1c) hosts and has been found in many Coleoptera, Diptera and Lepidoptera (Kageyama *et al*., 2012).

**Feminizing** (FE, Figure 1b) endosymbionts turn genetically male offspring into functional females by interfering with host sex differentiation pathways (Narita *et al*., 2007, Herran *et al*., 2021). This phenotype has so far been only defined in diplodiploid hosts, for example in butterflies and woodlice (Kageyama *et al*., 2012).

**Parthenogenesis induction** (PI, Figure 1d) occurs in haplodiploid hosts, for example in many Hymenoptera, where unfertilized eggs normally develop as males (arrhenotoky). Infected females switch from arrhenotoky to thelytoky, i.e., their unfertilized eggs develop as females (Ma & Schwander, 2017). The term ‘parthenogenesis induction’ is somewhat unfortunate for a switch from arrhenotokous to thelytokous parthenogenesis (Klein *et al*., submitted manuscript, shows the relevance of considering arrhenotoky a form of parthenogenesis too). However, we will stick to the term PI to avoid deviating from established terminology.

**Distortion of sex allocation** (SD, Figure 1e) likewise only occurs in haplodiploids, for example in the mite *Tetranychus urticae* (Wybouw *et al*., 2023). SD symbionts bias maternal resource investment towards an elevated fertilization rate (Vala *et al*., 2003; Wang *et al*., 2020; Bagheri *et al*., 2022), thereby increasing female production. One mechanism involves egg size increase (Katlav *et al*., 2022; Wybouw *et al*., 2023), since in haplodiploids, larger eggs are more likely to be fertilized (Macke *et al*., 2011; Katlav *et al*., 2021). Interestingly, SD can occur either as an independent manipulation (Vala *et al*., 2003; Wang *et al*., 2020), or together with CI through infected males (Katlav *et al*., 2022; Wybouw *et al*., 2023) — a mechanism where paternal infection status matters, as we explain in detail below.

### (2) Paternal-origin manipulation

All known examples of manipulations that depend on paternal infection status are forms of **cytoplasmic incompatibility** (CI, Figure 1f-h). CI is arguably the most widespread type of reproductive manipulation and occurs both in diplodiploid and haplodiploid hosts (Kageyama *et al*., 2012; Turelli *et al*., 2022). While the term CI is used on its own for diplodiploid hosts (Figure 1f), haplodiploids necessitate defining two subcategories (Figure 1g-h). Across all cases, the manipulation occurs if an infected male mates with a female not infected with the same strain (‘incompatible cross’).

In the incompatible cross, the mother is uninfected and so are all her offspring. It would now be counterproductive for the parasite to increase female progeny among those offspring (the adaptive response in the maternal-origin manipulations). Being stuck in a dead-end (the male), all the symbiont can do is to ‘sabotage’ the reproduction of the female: her uninfected progeny – should they develop normally – form competitors of infected females in the next generation. The three CI manifestations differ in whether the manipulation additionally decreases, causes no change, or increases the production of uninfected males.

In diplodiploid hosts, offspring from incompatible crosses die (or have starkly reduced viability) regardless of offspring sex (Figure 1f). Infected females produce viable offspring regardless of the infection status of their mating partner — provided that both sexes carry the same strain in case both are infected. Mechanistically, a ‘modification and rescue’ model (Werren, 1997; Hochstrasser, 2023) operates where CI-symbionts modify the sperm of infected males to reduce offspring viability, but the symbiont’s presence in the female counteracts (‘rescues’) this modification.

Under haplodiploidy, fertilization leads to diploid females; males are produced parthenogenetically. Since CI can only occur if there is fertilization, it can only affect diploid progeny, which, without CI, would develop as females. Depending on the extent of CI-induced defects in sperm (Serbus *et al*., 2008), the would-be females face two different fates, hence the two different CI types in haplodiploids: ‘**Female Mortality CI**’ (CI_FM_), where female progeny from an incompatible cross die (Figure 1g) and ‘**Male Development CI**’ (CI_MD_), where they develop into haploid males (Figure 1h).

## III. Features of the minimal model

We present a minimal model structure for all types of reproductive manipulation, with the goal of integrating across specific cases (and combinations thereof) that have been explored in previous work (Table 1). Infection dynamics are strongly dependent on three factors: (i) efficiency of vertical transmission (i.e. the proportion of eggs that are infected, given that the mother is infected), (ii) efficiency of the reproductive manipulation, and (iii) additional fitness effects conferred by the endosymbiont, unrelated to the reproductive manipulation. For the minimal model, transmission efficiency, denoted *T*, and the efficiency of reproductive manipulation, *β*, are obviously important.

**Table 1:** Existing theoretical studies of the infection dynamics of reproductive manipulators. Grey boxes represent combinations that are biologically unlikely because of host ploidy differences.

| invading symbiont →<br>resident population ↓ | MK | FE | CI | PI | SD |
| --- | --- | --- | --- | --- | --- |
| uninfected | Hurst, 1991b<br>Randerson *et al.*, 2000b<br>Groenenboom &<br>Hoegeweg 2002<br>Dyer & Jaenike, 2004<br>Zug & Hammerstein 2018 | Hatcher & Dunn, 1995<br>Randerson *et al.*, 2001 | Caspari & Watson, 1959<br>Fine, 1978<br>Turelli, 1994<br>Hoffmann *et al.*, 1990<br>Vavre *et al.*, 2000<br>Randerson *et al.*, 2001<br>Egas *et al.*, 2002<br>Hurst *et al.*, 2002<br>Keeling *et al.*, 2003<br>Flor *et al.*, 2007<br>Engelstädter & Telschow,<br>2009<br>Zug & Hammerstein, 2018<br>Karisto *et al.*, 2022 | Vavre *et al.*, 2000 | Vavre *et al.*, 2000*<br>Egas *et al.*, 2002*<br>Wybouw *et al.*, 2023 |
| MK | Randerson *et al.*, 2000b<br>Groenenboom &<br>Hoegeweg, 2002 |  | Engelstädter *et al.*, 2004; 2007<br>Zug & Hammerstein, 2018 |  |  |
| FE |  |  | Randerson *et al.*, 2001 | biologically unlikely | biologically unlikely |
| CI | Hurst *et al.*, 2002<br>Engelstädter *et al.*, 2004;<br>2007<br>Zug & Hammerstein, 2018 | Randerson *et al.*, 2001<br>Hurst *et al.*, 2002 | Frank, 1998<br>Turelli, 1994<br>Keeling *et al.*, 2003 | Hurst *et al.*, 2002<br>Engelstädter *et al.*,<br>2004 |  |
| PI |  | biologically unlikely | Engelstädter *et al.*, 2004 |  |  |
| SD |  | biologically unlikely |  |  |  |

\*Authors consider variable sex ratios in their equations for CI symbionts, which is somewhat equivalent of the SD case, although back then they did not explicitly know of it

Symbiont spread is compromised if infected females can produce uninfected eggs (*T* < 1), and recorded transmission efficiencies for endosymbionts are often high (Jiggins *et al*., 2002; Charlat *et al*., 2009). A failure to manipulate host reproduction (*β* < 1) makes parasites end up in dead-end hosts (males) or otherwise fail to use the mechanism that enhances their spread. Finally, given that transmission is vertical, it should pay for symbionts to additionally increase host fitness, modelled as a multiplicative effect on fitness, denoted *F* (Brownlie *et al*., 2009; Nikoh *et al*., 2014). This creates an interesting situation for the host, as benefits of being infected (*F* > 1) can coexist with significant costs due to sex ratio distortion, which in some cases involves a significant proportion of offspring dying (MK).

For all cases that fit into our framework, we derive invasion conditions and equilibrium frequencies for symbiont spread from rare, both in uninfected populations and in populations already infected with a reproductive manipulator of the same or a different type. For cases where there is a minimum frequency that must be reached before the frequency can increase further (i.e., CI), we also derive this threshold frequency. Some studies make useful deviations from the core assumptions that are not easily captured by our minimum model structure. We discuss these deviations separately.

### (1) Key assumptions

We consider a panmictic host population with non-overlapping generations. Uninfected females are assumed to produce an unbiased sex ratio. There are up to two infections in the population: either two reproductive manipulators of the same type (e.g., two male-killers, with potentially unequal fitness effects) or two different types (e.g., one male-killer and one CI-endosymbiont). We exclude coinfection at the individual host level, by assuming exclusively vertical transmission and an in initial host population that lacks coinfected individuals. Thus, any given population comprises up to six different categories of individuals: (two sexes) × (infected by one or the other symbiont, or uninfected). Hosts are either diplodiploid or haplodiploid (Figure 1).

Each strain is assumed to induce one type of reproductive manipulation (MK, FE, PI, SD, or CI). We examine the effects of vertical transmission efficiency *T*, reproductive manipulation efficiency *β*, and increases or decreases of female host fitness (other than the actual reproductive manipulation) by *F* > 1 and *F* < 1, respectively (Table 2 summarizes all symbols). *F* is defined as the fecundity of an infected female relative to an uninfected one (Zug & Hammerstein, 2018; Karisto *et al*., 2022).

**Table 2:**
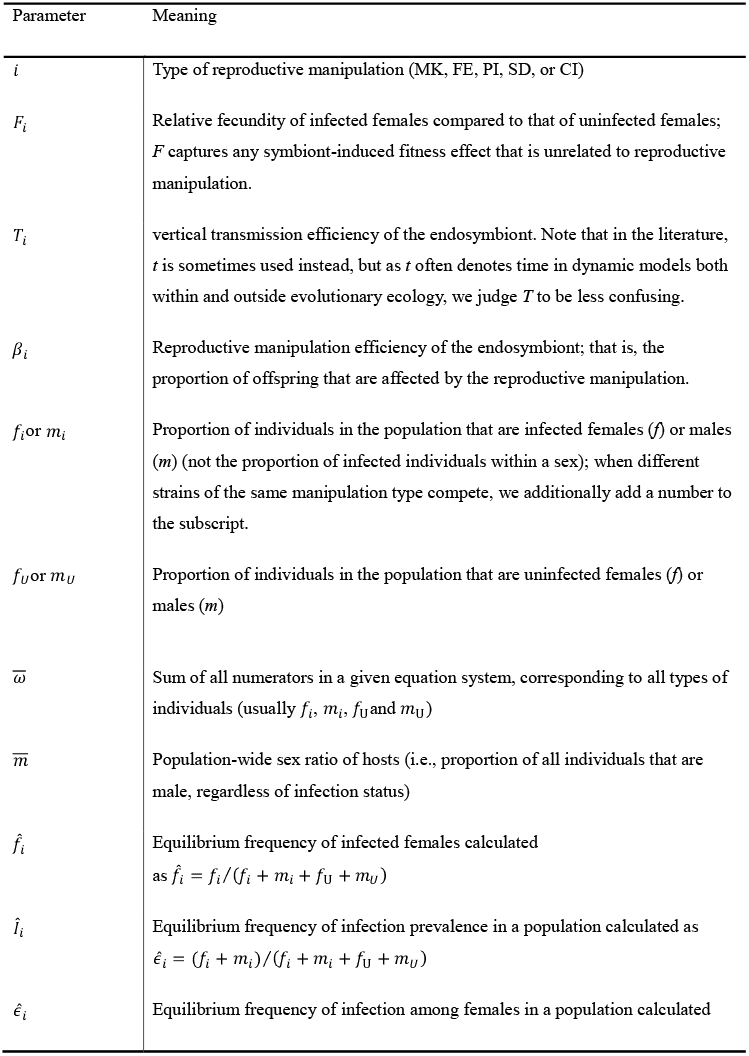
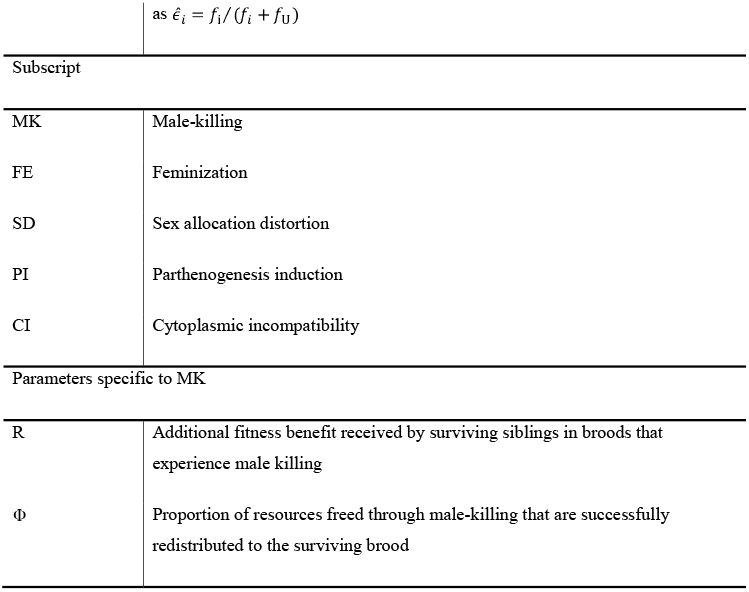
Symbols and meanings.

Importantly, *F* only captures fitness effects that are not related to reproductive manipulation: e.g., male offspring dying due to MK does not reduce *F* but is accounted for via *β* instead. In line with earlier theory, we also assume that *F* has strictly sex-limited effects, impacting females only (but see section VII.5).

Often, *F, β* and *T* operate multiplicatively (see Figure 2a for a visual example in the case of FE). The product *FT*, the ‘effective fecundity’ of an infected female (from the endosymbiont’s point of view), plays a conceptually important role in the infection dynamics (Turelli, 1994; Zug & Hammerstein, 2018; Karisto *et al*., 2022). The effects of *β* differ between manipulation types: for example, in a MK scenario *β* is interpreted as the probability that an infected egg, conditional on being male, dies, while in a CI_MD_ scenario *β* denotes the probability that an egg resulting from a cross between uninfected female and CI-infected male develops as a male.

**Figure 2:**
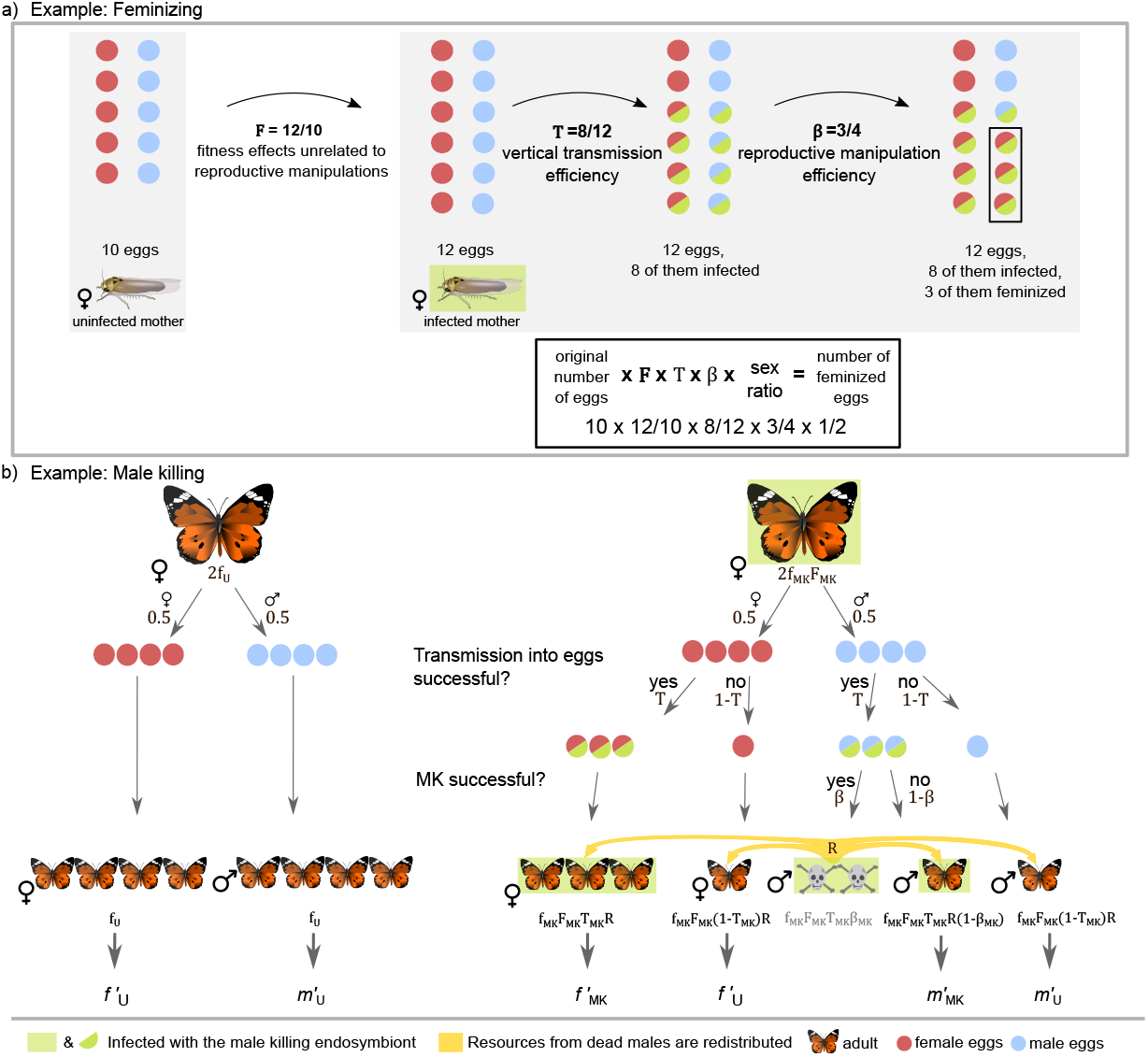
Schematic visualization of model parameters, using the example of feminization (a), and of a decision tree, using the example of male killing (b). The MK decision tree corresponds to equations 2.1-2.4. Eggs with green edges and individuals on a green background are infected, red eggs are female, blue eggs are male.

## IV. Infection dynamics: the case of a single strain

We begin by describing dynamics with just one strain, tracking the frequencies of uninfected females *f*_U_ and males *m*_U_, as well as infected females *f*_*i*_ and males *m*_*i*_ (where *i* stands for MK, FE, …, CI), from one generation to the next.

Exclusive vertical transmission means that infected offspring must have infected mothers, while uninfected offspring may have uninfected or infected mothers (under imperfect transmission efficiency *T*<1). This creates ‘leakage’ from infected to uninfected individuals unless *T* = 1. An endosymbiont will only spread if it makes up for this leakage, using any of the mechanisms described above.

In our first example (MK), we discuss the derivation of each equation in detail and then follow the same rationale for all subsequent manipulations, while also pointing out differences between manipulation types.

### (1) Male-killing (MK)

MK presents a conundrum: given that males are a dead-end for maternally transmitted symbionts, what is the benefit for the endosymbiont to die with the male egg, rather than later with the adult male? If the death of the male allows some reallocation of resources to female siblings (which are, to a large degree, infected with the same symbiotic strain), then the MK phenotype will aid the parasite’s spread. In other words, the male-killer boosts the fitness of female offspring at the expense of males, provided that infected females profit from the resources freed by the death of their brothers.

This is why models of MK dynamics (Hurst, 1991*b*; Randerson *et al*., 2000*b*; Zug & Hammerstein, 2018; Fisher *et al*., 2022) start by assuming that siblings in a brood compete for resources. The standard assumption is that resources are divided equally among all siblings and that individual fitness is proportional to the amount of acquired resources. When the mother is infected with a male-killer, a proportion *β*_MK_*T*_MK_/2 of her offspring die (half are males, and of those, a proportion *T*_MK_ inherits the male-killer, which then successfully kills with probability *β*_MK_). The resources freed by the death of infected males — a proportion *β*_MK_*T*_MK_/2 of all offspring resources — are available for redistribution among the survivors, which constitute a proportion 1–*β*_MK_*T*_MK_/2 of the original brood. We denote by *Φ* the proportion of freed resources that are successfully transferred to the surviving brood; the rest remains unused. Hence, each survivor obtains a resource boost of magnitude 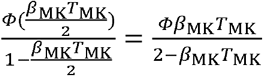. Assuming a linear relationship between resources and individual fitness (a somewhat strong assumption), the fitness of the surviving offspring is

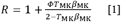

relative to the fitness of offspring of uninfected mothers, which is set to 1. We now have everything at hand to present the dynamics from one generation to the next. Recall that a proportion *β*_MK_ of infected male offspring die, leading to elevated fitness *R* > 1 for all remaining offspring of infected females. A proportion 1 − T_MK_ of female offspring of infected females do not inherit the endosymbiont and count towards the number of future uninfected females *f* ′_U_. An equivalent statement holds for male offspring. It follows that the next-generation frequencies of each class of host individual (see Figure 2b for a decision tree) obey

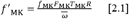

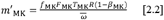

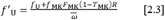

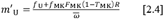

To move from counts to frequencies, the total production of each type of individual (numerators of eq. 2.1-2.4) is divided by the total production of all types, 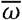, which is the sum of all numerators:

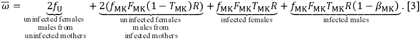

With *f*_MK_ = 0, 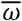 simplifies to 2*f*_U_ = 1, as uninfected females in that case form half of the population, the rest being uninfected males. We use this result when following the approach of Kriesner *et al*., (2013) and Zug & Hammerstein (2018) to investigate the fate of a rare male-killer in an otherwise uninfected population. The invasion condition for the male-killer spread is *f*′_MK_ > *f*_MK_. From equation 2.1, the criterion takes the form 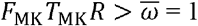, thus

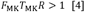

Summing up, the male-killer can increase in frequency if the product of its effective fecundity and the MK-associated fitness benefits (left-hand side of equation 4) is larger than 1, the fitness of uninfected females.

If invasion is successful, the frequency of male-killer infected females will rise until it reaches an equilibrium, *f*′_MK_ = *f*_MK_,, and we denote this equilibrium frequency as 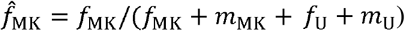,. Note that this equation describes the equilibrium frequency of infected females in the population. But there are other ways to indicate the equilibrium frequency of the infection, such as the overall infection frequency (i.e., infected males and females) in the population, *Î*_MK_ = (*f*_MK_ + *m*_MK_) / (*f*_MK_ + *m*_MK_ + *f*_U_ + *m*_U_), or the equilibrium frequency of infected females among females only,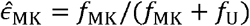. While most theoretical work denotes equilibria in terms of infected females in the population 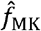 or among females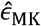, empirical work often records the overall infection prevalence *Î*_MK_. For this reason, our main plots (Figure 3) are based on overall prevalence, while we also provide equations for the other two, and provide plots that use these measures (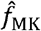 in Figure 4 & S7, and 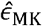 in Figure S9).

**Figure 3:**
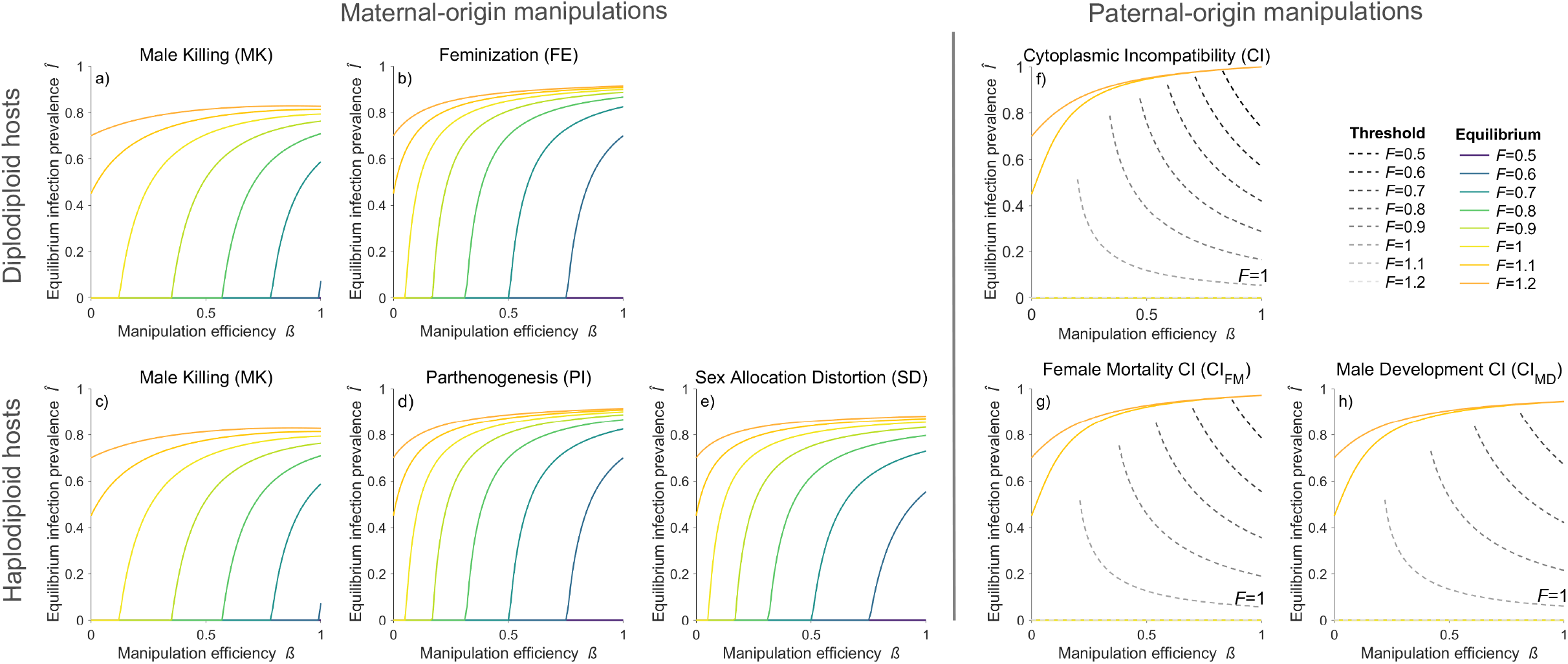
Equilibrium frequencies (overall infection prevalence) *Î* (coloured lines) and CI invasion thresholds 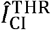 (grey dashed lines) for the different manipulation types, for varying manipulation efficiency *β* (x axis) and relative fecundity *F* (colour). Examples were derived with *T* 0.95, and, for MK, *Φ* 0.85. a), b) and f) show results for diplodiploid hosts, and c), d), e), g) and h) for haplodiploid hosts. For all parameter combinations that do not fulfil the CI invasion condition (equations 16, 19 and 22), invasion is not possible and hence the equilibrium frequency is zero, which in these examples applies to all cases where *F* <= 1 (f, g, h). For these cases, the corresponding threshold frequencies are shown (dashed lines). In contrast, for cases where *F* > 1, there is no invasion threshold, i.e., CI symbionts can invade from rarity and will reach the corresponding equilibrium frequency.

**Figure 4:**
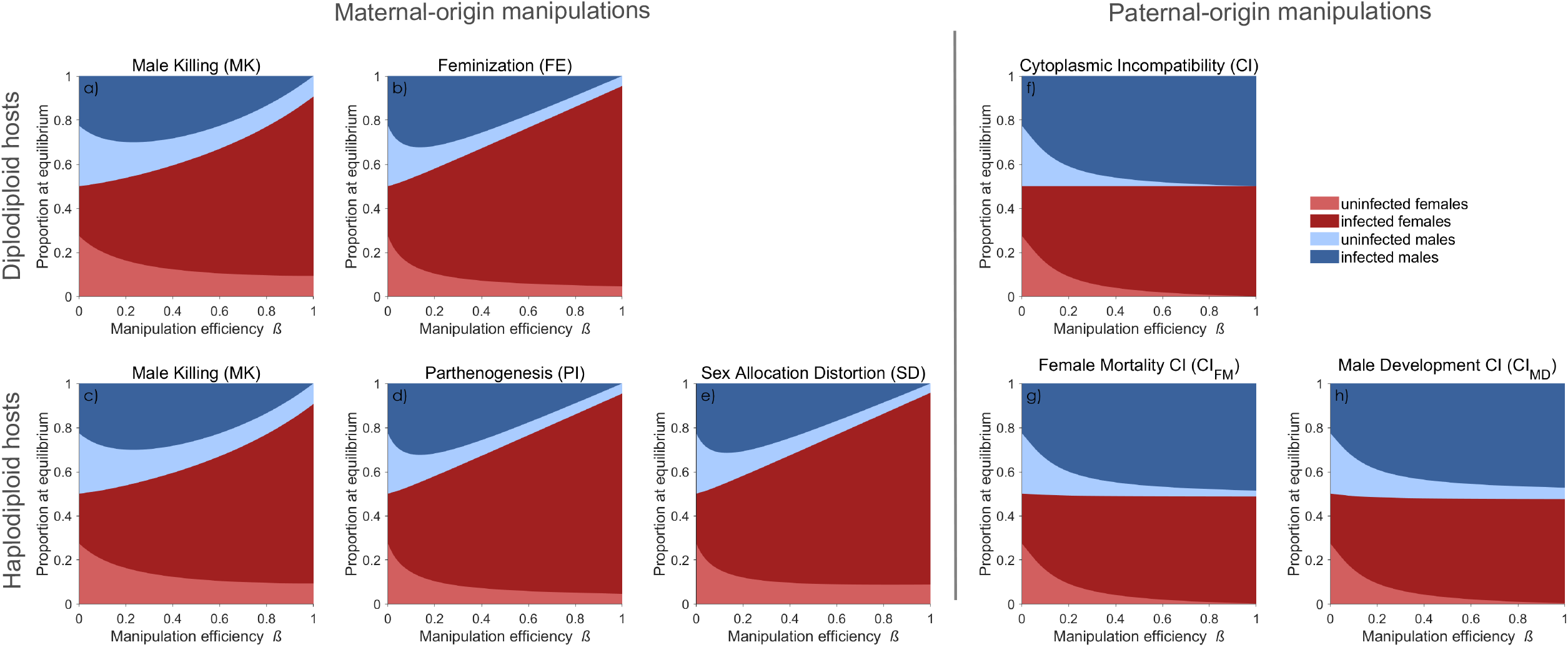
Proportion at equilibrium of all four types of individuals in the population (infected females, 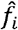; uninfected females,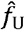; infected males,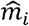; uninfected males,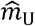) for the different manipulation types, for varying manipulation efficiency *β* (x axis). Examples were derived with *T* 0.95, *F* 1.1 and, for MK, *Φ* 0.85. a), b) and f) show results for diplodiploid hosts, and c), d), e), g) and h) for haplodiploid hosts.

Solving the above yields

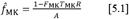

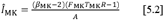

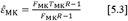

where *A* = 2 − *β*_MK_ − *F*_MK_ *R*(2+ *β*_MK_*T*_MK._

Equations 5.1-3 predicts high equilibrium frequencies when *β*_MK_ is high, especially if combined with high effective fecundity *F*_MK_*T*_MK_ (Figure 3a exemplifies the effect by showing the effect of *F*; varying *T* yields equivalent results due to the multiplicative nature of effective fecundity).

We are aware of only one case of MK in a haplodiploid species, the parasitoid wasp *Nasonia vitripennis* (Werren *et al*., 1986). MK infection dynamics do not differ between haplodiploid and diplodiploid hosts (Figure 3&4a, c, see also section VI.1).

Note that we have assumed that females are not sperm-limited, although the spread of a successful male-killer will cause population-wide male shortage. For this reason, some models of MK infection dynamics include spatial structure (Groenenboom & Hoegeweg, 2002; Bonte *et al*., 2008; 2009): if some subpopulations run out of males before others, immigrant males may help, and if this influx of males keeps the local dynamics away from local extinction, it may also allow infected females to spread the infection to neighbouring demes (see also section VII.3).

### (2) Feminization (FE)

Following previous work (Randerson *et al*., 2001), we assume that a proportion *β*_FE_ of infected male offspring are successfully feminized; the remaining ones develop as males. We also assume that *F* and *T* do not differ between genetic females and feminized males. Therefore, the next-generation frequencies of all types of individuals in the population (*f*_FE_,*m*_FE_,*f*_U_,*m*_U_) are

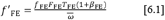

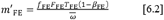

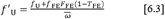

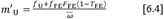

The feminized males are added to the pool of infected females (equation 6.1) and removed from equation 6.2 (Figure S1). A rare feminiser will successfully invade an uninfected population If

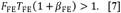

Thus, to invade, an infected mother must produce a sum of infected genetic females (*F*_FE_*T*_FE_) and successfully feminized males (*F*_FE_*T*_FE_*β*_FE_) that exceeds 1. Even if infection reduces effective fecundity, a sufficiently high success in feminizing male offspring can overcome this detrimental effect and make the symbiont invade. After successful invasion, the feminiser reaches the nontrivial equilibrium

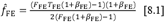

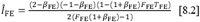

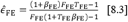

Compared with MK, FE appears a superior manipulation strategy for the parasite: symbionts in killed males are simply lost while they directly contribute to the next generation if feminized.

Correspondingly, positive equilibrium frequencies arise at far less perfect *β*_FE_ values than what is required for *β*_MK_ (compare Figure 3a-b or Figure S7a-b, though the exact comparison also depends on the MK-specific parameter value *Φ*, which we assumed to be moderately good, *Φ* = 0.85).

Despite these differences between MK and FE, they share a similarity: both have the potential to make a local host population run out of males. In FE, the problem appears more severe because (for otherwise similar parameter values) the equilibrium frequency of the parasite is higher (compare Figure 3a, b).

### (3) Parthenogenesis Induction (PI)

Given that PI has been reliably documented only in haplodiploid species, we assume host haplodiploidy. A proportion *β* _PI_ of infected unfertilized haploid eggs are assumed to develop as diploid female offspring. We further assume that infected eggs in which PI fails develop normally; that is, they can be fertilized and develop as females (we assume 50% probability for this), while the remaining 50% develop as males. This leads to the following recursion equations:

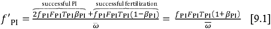

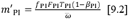

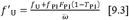

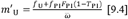

As before, 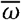 is the sum of all numerators (for visual derivation see Figure S2).

Note that we assume that PI, if successful, takes place before fertilization, and PI-transformed eggs can no longer be fertilized. This creates two routes to becoming an infected female: (1) successful PI (first term in the numerator of equation 9.1) and (2) successful fertilization after PI has failed (second term in the numerator of equation 9.1). Infected males are produced when PI was present in the mother but failed, and fertilization also did not occur (equation 9.2).

The condition for a rare PI-endosymbiont to invade into an uninfected population is given by

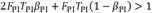

which simplifies to

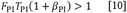

This equation takes the same form as the FE case did, and so does the equilibrium frequency

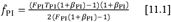

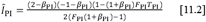

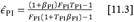

Thus, the equilibrium frequencies of PI in haplodiploids are identical to those of FE in diplodiploids for the same values of effective fecundity and *β* (Figure 3 & 4b, d) (see also section VI.3).

Once PI is common, male shortage is not a problem for a population since infected females can produce daughters parthenogenetically. This contrasts with MK and FE, where sperm limitation can become a serious issue should the symbiont approach fixation. Accordingly, in several PI-infected species, transmission efficiency and PI strength are nearly perfect, and infection has gone to fixation. The resulting all-female populations reproduce asexually (Ma & Schwander, 2017) and the term that accounts for successful fertilization in equations 9.1 and 10 is no longer needed.

### (4) Sex Allocation Distortion (SD)

Recent modelling of SD infection dynamics (Wybouw *et al*., 2023) builds upon CI models that considered the possibility of variable fertilization rates in infected haplodiploids (Vavre *et al*., 2000) and different sex ratios produced by uninfected and infected females (Egas *et al*., 2002) before SD was known from nature. Here, we model SD as a distinct manipulation type. We define the parameter *β*_SD_ as a shift from the original sex ratio towards females; if *β*_SD_ = 1, all offspring become female, while *β*_SD_ = 1 retains the original sex ratio of 1:1.

The recursion equations (Figure S3 offers a decision tree) are

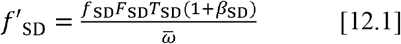

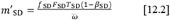

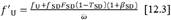

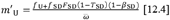

As usual, 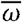 sums the numerators (equation 12.1-12.4).

The numerator of equations 12.3 and 12.4 is formed differently than for FE (6.3-4) and PI (9.3-4) because, with SD, we assume the brood’s sex ratio is impacted by the endosymbiont inhabiting the mother, not of the offspring. SD thus impacts offspring sex even if they are not themselves infected. Nevertheless, the invasion condition for SD is identical to that of FE and PI (equation 7 and 10, respectively):

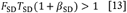

Consequently, SD, FE and PI all start to invade at the same parameter values of *F, T* and *β* (Figure 3b, d, e). The differences show at equilibria. While the equivalence of FE and PI extends to identical equilibrium frequencies, SD equilibrates at a lower frequency

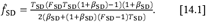

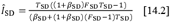

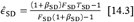

We assume that SD increases a mother’s infected and uninfected female offspring production, while FE or PI only increase the production of infected female offspring. This ‘dilution’ by uninfecteds makes SD equilibrate at a lower frequency (Figure 3e), a difference that is clearest at low transmission efficiency (Figure S7 & S8).

### (5) Cytoplasmic incompatibility (CI): the diplodiploid case

Thus far (IV.1-4) we considered maternal-origin manipulations. All cases of CI represent manipulations of paternal origin, which complicates the infection dynamics considerably. CI differs from our previous examples as it does not boost the production of infected females, but damages the fecundity of uninfected females, should they mate with infected males. The symbiont only receives a relative fitness boost. The intricate nature of this effect has created a considerable body of literature on CI infection dynamics (Caspari & Watson, 1959; Fine, 1978; Hoffmann *et al*., 1990; Turelli, 1994; Engelstädter & Telschow, 2009; Karisto *et al*., 2022).

Incompatible crosses between an infected male and a female that is not infected with the same strain result in the death of a proportion *β*_CI_ of offspring. Following the notation from Zug & Hammerstein (2018), we use round and square brackets to denote maternal and paternal contributions, respectively, to the next generation (Figure S4):

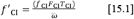

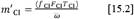

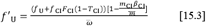

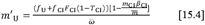

with 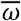 being the sum of all numerators.

Maternal contributions are straightforward; in the following, we describe how paternal contributions are derived. Assuming random mating, uninfected males sire a proportion 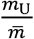 of offspring, while infected males sire a proportion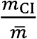 with 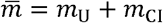. Infected females can mate with either type of male without suffering from CI; therefore, the paternal contribution in equations 15.1-2 is 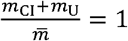 (hence no square brackets in either equation). Uninfected females (equations 15.3-4) experience CI if fathers are infected, in which case a proportion *β*_CI_ of the offspring will die. Here, the paternal contribution modifies offspring production by 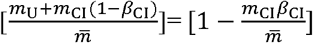.

The invasion condition is

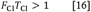

Strikingly, CI differs from all other reproductive manipulations in that its invasion success only depends on effective fecundity *F*_CI_*T*_CI_, not on CI efficiency *β*_CI_. This is in line with theory on selection on CI-symbionts, which acts on effective fecundity, whereas CI efficiency is selectively neutral (diplodiploid case: Prout, 1994; Turelli, 1994; haplodiploid case: Egas *et al*., 2002; Vavre *et al*., 2003). The neutrality of CI strength may appear surprising, but since the sperm modification function underlying CI is only expressed in males, it cannot be selected in maternally inherited symbionts.

Host population structure could reward good manipulators in principle (Hurst, 1991*a*; Frank 1997), but even here, selection remains weak and transient (Haygood & Turelli, 2009). Recent findings showing that genes involved in the modification function frequently acquire loss-of-function mutations and become pseudogenized (Martinez *et al*., 2021) align with the neutrality prediction.

The fact that CI invasion success does not depend on *β*_CI_ is a direct consequence of the inefficiency of CI when rare, rendering the frequency of incompatible matings very low. CI can only play to its strengths when sufficiently common. As discussed in section V.2, this positive frequency dependence represents a double-edged sword, as it prevents CI from invading when rare, but protects CI against invasion when common (assuming the invader is incompatible). Positive frequency dependence distinguishes CI from all other manipulation types and, if *F*_CI_*T*_CI_ < 1, introduces a threshold frequency that CI must reach to spread further (Caspari & Watson, 1959; Fine 1978). The dynamics involve three equilibria: the trivial equilibrium 0, an unstable equilibrium that gives the invasion threshold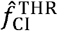, and a positive, stable equilibrium frequency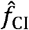. The latter two take the values

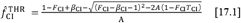

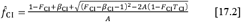

Since CI-infected females produce an equal sex ratio, the equilibrium frequency of infected individuals in the population is simply twice the frequency of infected females:

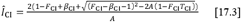

with *A* = 2*β*_CI_(1 − F_CI_ (+ *T*_CI_)).

The proportion of infected females among all females is equal to the proportion of infected individuals in the population, 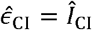, because both infected and uninfected individuals are produced at an equal sex ratio, in contrast to maternal-origin manipulations (e.g., MK, FE).

While high *β*_CI_ does not aid initial invasion, it strengthens the negative impact on uninfected females and thus lowers the invasion threshold (Figure 3f, grey dashed lines) as well as pushes CI to (slightly) higher equilibrium frequencies (Figure 3f, coloured lines). The invasion threshold also decreases if effective fecundity is high (comparison across dashed lines, Figure 3f). Since CI-infected females produce an equal sex ratio, frequencies of infected females 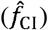 never exceed 0.5 (Figure S7).

However, this does not prevent infection prevalence from being high; prevalence is 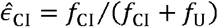 among females and *Î*_CI_ = (*f*_CI_ + *m*_CI_ / (*f*_CI_ + *m*_CI_ + *f*_U_ + *m*_U_), among all, and for these measures it has been a challenge to understand why low values are possible at all (Karisto *et al*. 2022).

### (6) Female-mortality CI in haplodiploids

In haplodiploid hosts, CI can only affect daughters, because male production does not require fertilization. With CI_FM_, a proportion 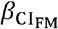 of daughters from incompatible crosses dies, leading to the following recursion equations (again, round and square brackets refer to maternal and paternal contributions, respectively; see Figure S5 for a decision tree):

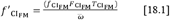

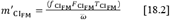

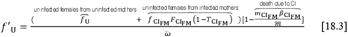

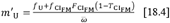

with 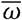 being the sum of all numerators, and 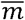 being the sum of all male frequencies in the population.

The invasion condition for CI_FM_ is

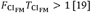

and the invasion threshold and the equilibrium frequencies are

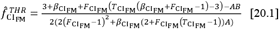

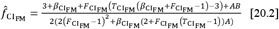

The equilibrium infection prevalence in the population and among females is

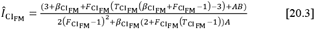

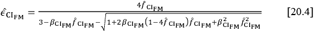

with 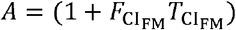 and 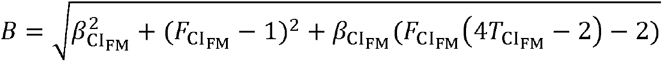.

Note that equations 20.4 and 23.4 involve 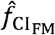 or 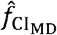 (as given by equations 20.2 and 23.2, respectively), which makes them less cumbersome. We discuss similarities and differences of the different CI types in diplodiploid and haplodiploid hosts in section VI.2.

### (7) Male-development CI in haplodiploids

With CI_MD_, a proportion 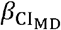 of daughters from incompatible crosses is turned into males, leading to the following recursion equations (see Figure S6 for the decision tree):

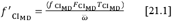

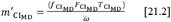

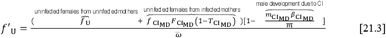

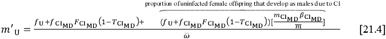

with 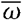 and 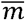 summing up all numerators and male-specific numerators, respectively. Uninfected males develop not only from unfertilized, uninfected eggs (first two terms in the numerator in equation 21.4), but also from incompatible crosses between uninfected females and infected males (third term in equation 21.4).

The invasion condition for CI_MD_, like that of all other CI cases, depends only on effective fecundity, without any effect of *β*:

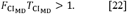

The threshold and equilibrium frequencies are, respectively:

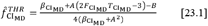

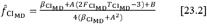

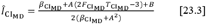

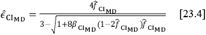

with 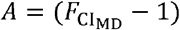 and 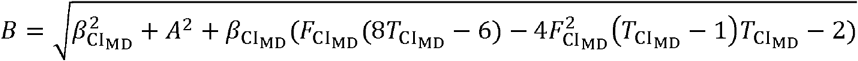.

Section VI.2 discusses similarities and differences of CI infection dynamics in diplodiploid vs. haplodiploid hosts.

## V. Infection dynamics involving two different types of reproductive manipulators

### (1) The basic rate of increase

Our above invasion conditions for a single symbiont strain invading into an uninfected host population all exhibit a general structure

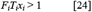

where the effective fecundity *F*_*i*_*T*_*i*_ of a strain *i* combines (if *x*_*i*_ > 1) with an additional boost when reproductive manipulation is successful. Usually, *x*_*i*_ increases with the efficiency of reproductive manipulation *β*_*i*_, (first row in Tables 3 and 4); but for CI this is not the case, as explained in section IV.5-7. This simple product combines transmission efficiency *T*_*i*_ with fitness effects either causally related to reproductive manipulation (*x*_*i*_) or not (*F*_*i*_). From the perspective of the maternally transmitted symbiont, the term *F*_*i*_*T*_*i*_*x*_*i*_ describes its ‘basic rate of increase’ (*sensu* Randerson *et al*., 2000*b*).

**Table 3:** Diplodiploids – Invasion Conditions.

|  |  |  |  |
| --- | --- | --- | --- |
| invading symbiont<br>→<br>resident<br>population ↓ | MK | CI | FE |
| uninfected | $F_{MK}T_{MK}R > 1$ | $F_{CI}T_{CI} > 1$ | $F_{FE}T_{FE}(1 + \beta_{FE}) > 1$ |
| MK | $F_{MK_2}T_{MK_2}R_{MK_2} > F_{MK_1}T_{MK_1}R_{MK_1}$ | $F_{CI}T_{CI} > F_{MK}T_{MK}R$ | $F_{FE}T_{FE}(1 + \beta_{FE}) > F_{MK}T_{MK}R$ |
| CI | $F_{MK}T_{MK}R \left[ 1 - \frac{m_{CI}\beta_{CI}}{\bar{m}} \right] > F_{CI}T_{CI}$ | $F_{CI_2}T_{CI_2} \left[ 1 - \frac{m_{CI_1}\beta_{CI_1}}{\bar{m}} \right] > F_{CI_1}T_{CI_1}$ | $F_{FE}T_{FE}(1 + \beta_{FE}) \left[ 1 - \frac{m_{CI}\beta_{CI}}{\bar{m}} \right] > F_{CI}T_{CI}$ |
| FE | $F_{\text{MK}} T_{\text{MK}} R > F_{\text{FE}} T_{\text{FE}} (1 + \beta_{\text{FE}})$ | $F_{\text{CI}} T_{\text{CI}} > F_{\text{FE}} T_{\text{FE}} (1 + \beta_{\text{FE}})$ | $F_{\text{FE}_2} T_{\text{FE}_2} (1 + \beta_{\text{FE}_2}) > F_{\text{FE}_1} T_{\text{FE}_1} (1 + \beta_{\text{FE}_1})$ |

**Table 4:** Haplodiploids – Invasion Conditions.

| invading<br>symbiont →<br>resident<br>population ↓ | CI<sub>FM</sub> | CI<sub>MD</sub> | MK | PI | SD |
| --- | --- | --- | --- | --- | --- |
| uninfected | $F_{\text{CI}_{\text{FM}}} T_{\text{CI}_{\text{FM}}} > 1$ | $F_{\text{CI}_{\text{MD}}} T_{\text{CI}_{\text{MD}}} > 1$ | $F_{\text{MK}} T_{\text{MK}} R > 1$ | $F_{\text{PI}} T_{\text{PI}} (1 + \beta_{\text{PI}}) > 1$ | $F_{\text{SD}} T_{\text{SD}} (1 + \beta_{\text{SD}}) > 1$ |
| CI<sub>FM</sub> | $F_{\text{CI}_{\text{FM}_2}} T_{\text{CI}_{\text{FM}_2}} \left[ 1 - \frac{m_{\text{CI}_{\text{FM}_1}} \beta_{\text{CI}_{\text{FM}_1}}}{\bar{m}} \right] > F_{\text{CI}_{\text{FM}_1}} T_{\text{CI}_{\text{FM}_1}}$ | $F_{\text{CI}_{\text{MD}}} T_{\text{CI}_{\text{MD}}} \left[ 1 - \frac{\beta_{\text{CI}_{\text{FM}}} m_{\text{CI}_{\text{FM}}}}{\bar{m}} \right] > F_{\text{CI}_{\text{FM}}} T_{\text{CI}_{\text{FM}}}$ | $F_{\text{MK}} T_{\text{MK}} R \left[ 1 - \frac{m_{\text{CI}_{\text{FM}}} \beta_{\text{CI}_{\text{FM}}}}{\bar{m}} \right] > F_{\text{CI}_{\text{FM}}} T_{\text{CI}_{\text{FM}}}$ | $2F_{\text{PI}} T_{\text{PI}} \beta_{\text{PI}} + (F_{\text{PI}} T_{\text{PI}} (1 - \beta_{\text{PI}}) [1 - \frac{m_{\text{CI}_{\text{FM}}} \beta_{\text{CI}_{\text{FM}}}}{\bar{m}}]) > F_{\text{CI}_{\text{FM}}} T_{\text{CI}_{\text{FM}}}$ | $F_{\text{SD}} T_{\text{SD}} (1 + \beta_{\text{SD}}) [1 - \frac{m_{\text{CI}_{\text{FM}}} \beta_{\text{CI}_{\text{FM}}}}{\bar{m}}] > F_{\text{CI}_{\text{FM}}} T_{\text{CI}_{\text{FM}}}$ |
| CI<sub>MD</sub> | $F_{\text{CI}_{\text{FM}}} T_{\text{CI}_{\text{FM}}} \left[ 1 - \frac{\beta_{\text{CI}_{\text{MD}}} m_{\text{CI}_{\text{MD}}}}{\bar{m}} \right] > F_{\text{CI}_{\text{MD}}} T_{\text{CI}_{\text{MD}}}$ | $F_{\text{CI}_{\text{MD}_2}} T_{\text{CI}_{\text{MD}_2}} \left[ 1 - \frac{m_{\text{CI}_{\text{MD}_1}} \beta_{\text{CI}_{\text{MD}_1}}}{\bar{m}} \right] > F_{\text{CI}_{\text{MD}_1}} T_{\text{CI}_{\text{MD}_1}}$ | $F_{\text{MK}} T_{\text{MK}} R \left[ 1 - \frac{m_{\text{CI}_{\text{MD}}} \beta_{\text{CI}_{\text{MD}}}}{\bar{m}} \right] > F_{\text{CI}_{\text{MD}}} T_{\text{CI}_{\text{MD}}}$ | $2F_{\text{PI}} T_{\text{PI}} \beta_{\text{PI}} + (F_{\text{PI}} T_{\text{PI}} (1 - \beta_{\text{PI}}) [1 - \frac{m_{\text{CI}_{\text{MD}}} \beta_{\text{CI}_{\text{MD}}}}{\bar{m}}]) > F_{\text{CI}_{\text{MD}}} T_{\text{CI}_{\text{MD}}}$ | $F_{\text{SD}} T_{\text{SD}} (1 + \beta_{\text{SD}}) [1 - \frac{m_{\text{CI}_{\text{MD}}} \beta_{\text{CI}_{\text{MD}}}}{\bar{m}}] > F_{\text{CI}_{\text{MD}}} T_{\text{CI}_{\text{MD}}}$ |
| MK | $F_{\text{CI}_{\text{FM}}} T_{\text{CI}_{\text{FM}}} > F_{\text{MK}} T_{\text{MK}} R$ | $F_{\text{CI}_{\text{MD}}} T_{\text{CI}_{\text{MD}}} > F_{\text{MK}} T_{\text{MK}} R$ | $F_{\text{MK}_2} T_{\text{MK}_2} R_{\text{MK}_2} > F_{\text{MK}_1} T_{\text{MK}_1} R_{\text{MK}_1}$ | $F_{\text{PI}} T_{\text{PI}} (1 + \beta_{\text{PI}}) > F_{\text{MK}} T_{\text{MK}} R$ | $F_{\text{SD}} T_{\text{SD}} (1 + \beta_{\text{SD}}) > F_{\text{MK}} T_{\text{MK}} R$ |
| **PI** | $F_{CI_{FM}} T_{CI_{FM}} > F_{PI} T_{PI} (1 + \beta_{PI})^*$ | $F_{CI_{MD}} T_{CI_{MD}} > F_{PI} T_{PI} (1 + \beta_{PI})^*$ | $F_{MK} T_{MK} R > F_{PI} T_{PI} (1 + \beta_{PI})^*$ | $F_{PI_2} T_{PI_2} (1 + \beta_{PI_2}) > F_{PI_1} T_{PI_1} (1 + \beta_{PI_1})^{**}$ | $F_{SD} T_{SD} (1 + \beta_{SD}) > F_{PI} T_{PI} (1 + \beta_{PI})^*$ |
| **SD** | $F_{CI_{FM}} T_{CI_{FM}} > F_{SD} T_{SD} (1 + \beta_{SD})$ | $F_{CI_{MD}} T_{CI_{MD}} > F_{SD} T_{SD} (1 + \beta_{SD})$ | $F_{MK} T_{MK} R > F_{SD} T_{SD} (1 + \beta_{SD})$ | $F_{PI} T_{PI} (1 + \beta_{PI}) > F_{SD} T_{SD} (1 + \beta_{SD})$ | $F_{SD_2} T_{SD_2} (1 + \beta_{SD_2}) > F_{SD_1} T_{SD_1} (1 + \beta_{SD_1})$ |

\* Here we assume that $T_{PI} < 1$ and $\beta_{PI} < 1$ , which means that there are males in the resident population.
\*\* Here we assume that $T_{PI_2} < 1$ and $\beta_{PI_2} < 1$ , which means that there are males in the resident population. If $T_{PI_2} = 1$ and $\beta_{PI_2} = 1$ , there are no males in the resident population and
the invasion condition boils down to: $2F_{PI_2} T_{PI_2} \beta_{PI_2} > 2F_{PI_1} T_{PI_1} \beta_{PI_1}$

We can now further generalize the structure of the invasion condition to include invasion scenarios where the resident population is infected at equilibrium with a different strain or an entirely different type of reproductive manipulator. In such two-strain (e.g., MK_1_, MK_2_) or two-type (e.g., CI, MK) scenarios, the invasion condition exhibits the following general structure:

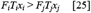

where the subscripts *i* and *j* represent two different strains or types. As an example, consider the case of a feminizer invading into a population infected with a male-killer. The basic rate of increase *F*_FE_*T*_FE_(1+*β*_FE_) of the feminizer is simply compared with that of the male-killer, *F*_MK_*T*_MK_*R*, and the invasion condition is

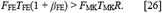

In sum, the strain with the higher basic rate of increase will outcompete the other strain; stable coexistence is not possible. Our comparisons thus generalize the insight from previous theoretical work (two strains, CI: Caspari & Watson, 1959; Rousset *et al*., 1991; MK: Randerson *et al*., 2000*b*; two types: CI-MK, CI-PI: Engelstädter *et al*., 2004). However, this insight relies on the simplicity of our basic framework, and stable coexistence can become possible if assumptions are relaxed, e.g. by incorporating host population structure or coinfections within one host individual (see section VII).

### (2) Invasion into CI-infected populations: the priority effect

The above logic makes it straightforward to derive invasion conditions for most combinations of invading and resident strains. The exceptions that require more careful analysis are scenarios where the resident population is infected with CI. Here invading females do not carry the resident CI strain and suffer reduced fecundity in incompatible matings with resident CI-infected males (the proportion of surviving offspring is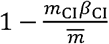, equations 15.3 and 15.4).

This makes CI well equipped to resist invasion by incompatible strains, that is, by strains that cannot rescue the sperm modification induced by the resident CI strain (Hurst, 1991*a*). This creates a priority effect (Zou & Rudolf, 2023) which protects the resident CI strain against invasion. The priority effect can be overcome if the invading strain is able to rescue the resident strain’s CI (Hurst *et al*., 2002) and/or if it exhibits a considerably larger effective fecundity than the resident strain (Zug & Hammerstein, 2018). Ultimately, this priority effect arises from CI’s positive frequency dependence (Zou & Rudolf, 2023). Hence, the double-edged sword of positive frequency dependence comes with an upside when CI is common (the priority effect) and a downside when CI is rare (the invasion threshold).

We will now briefly sketch two examples where the resident population has CI, one for diplodiploid and one for haplodiploid hosts. First, consider a diplodiploid scenario involving two different CI strains. The invasion condition is

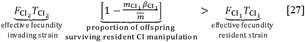

Note that we assume bidirectional CI, i.e., both CI strains are incompatible with each other. This means that the invading strain cannot rescue CI, hence there is a priority effect favouring the resident strain: due to its rarity, the invading strain imposes negligible negative effects on the resident strain, but experiences strong fitness loss induced by the resident strain.

For a second example, consider a PI-symbiont invading into a CI_FM_-infected population. The invasion condition follows a similar logic

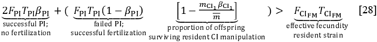

To invade a CI_FM_-infected population, PI endosymbionts need to overcome the detrimental effects of the resident CI strain. However, since PI enables the production of infected female offspring without mating (first term on the left-hand side), CI loses much of its manipulative power (now only affecting the second term on the left hand-side of equation 28). PI is thus predicted to invade CI-infected populations better than manipulation types (MK, FE, SD) that produce infected females via fertilized eggs.

The above information is sufficient to derive the invasion conditions for all possible two-strain and two-type combinations within our framework. Therefore, rather than presenting each case in detail, we refer the reader to Tables 3 and 4 for invasion conditions involving diplodiploid and haplodiploid hosts, respectively. The recursion equations for each invasion scenario are listed in Supplementary Tables S1 (diplodiploids) and S2 (haplodiploids).

## VI. Similarities and differences in infection dynamics in haplodiploid vs. diplodiploid hosts

Above, we structured our work based on comparing maternal-origin and paternal-origin manipulations. It is similarly useful to compare the other ‘axis’ of Figure 1, i.e. whether hosts are diplodiploid or haplodiploid. MK and CI are known from both types of hosts, FE is only known from diplodiploids, and PI and SD have only been convincingly documented in haplodiploids (Kageyama *et al*., 2012; Wybouw *et al*., 2023). For MK and CI, it is therefore possible to compare the infection dynamics across host ploidy systems and assess the relative ease or difficulty of invasion. It is also worthwhile to compare FE (in diplodiploids) with PI (in haplodiploids).

### (1) MK in diplodiploid vs. haplodiploid hosts

Although the mechanisms underlying MK differ between haplodiploid vs. diplodiploid hosts (Hornett *et al*., 2022), these create no difference for the infection dynamics (assuming equivalent parameter values) because the manipulation outcome, the death of males, does not depend on how the male-killer achieves its lethality, or on offspring ploidy. Then invasion conditions and equilibria are identical (compare Figure 3a, d) assuming that (1) sex ratios in uninfected hosts are balanced (regardless of whether they are diplodiploid or haplodiploid), (2) the death of infected brothers frees resources that can be used by female siblings in an equivalent manner.

### (2) CI in diplodiploid vs. haplodiploid hosts

All CI types damage the production of uninfected females; differences between CI types concern the production of uninfected males: a decrease in the case of CI in diplodiploids, no change with CI_FM_, and an increase with CI_MD_. In essence, the more uninfected offspring survive the CI manipulation, the harder it gets for CI to “do its job”. This leads to a clear trend from CI in diplodiploids over CI_FM_ to CI_MD_, which is characterized by a decrease in the equilibrium frequency (Figure 5a, b, c; unless *T* = 1; in this case, infection goes to fixation in all cases) and an increase in the threshold frequency (Figure 3f, g, h).

**Figure 5:**
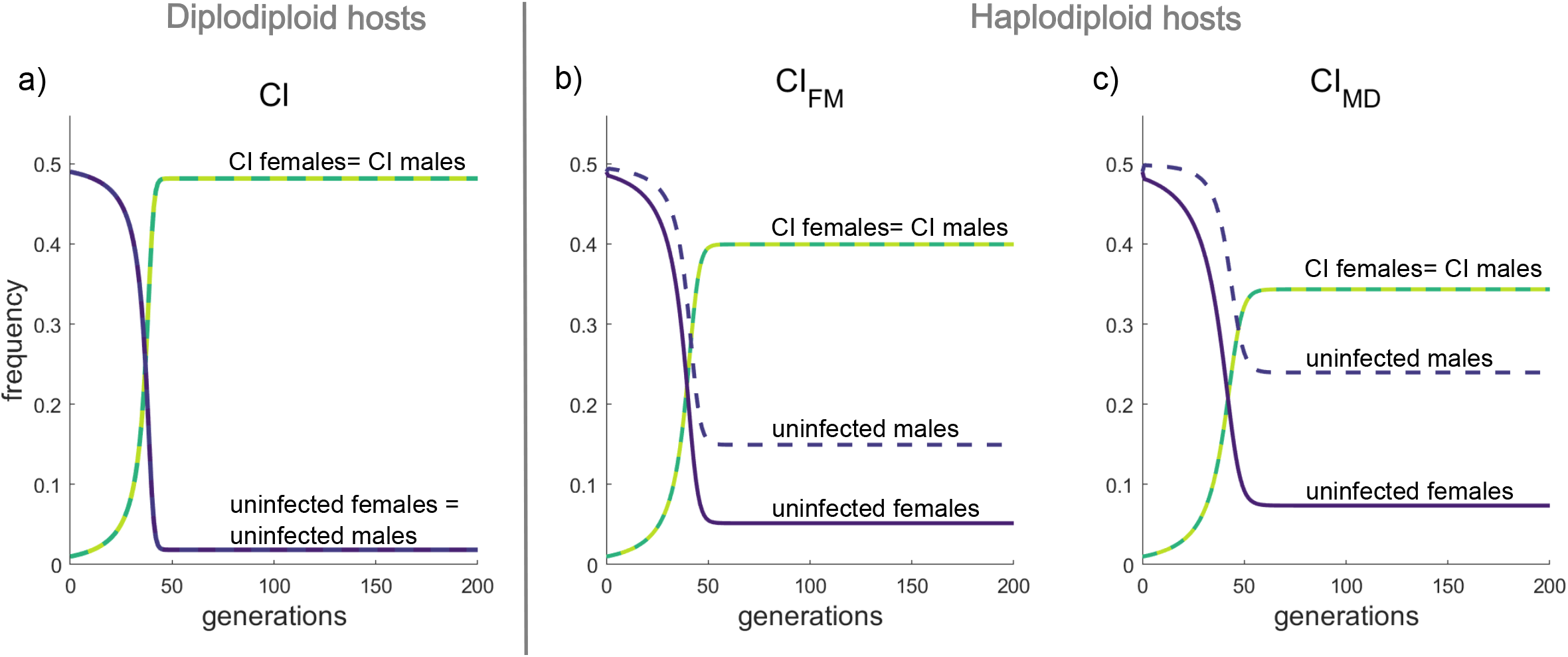
Similarities and differences in CI infection dynamics in diplodiploid (a) and haplodiploid hosts (b: CI_FM_; c: CI_MD_), showing the change of frequency of males *m*_U_, *m*_i_ (dashed lines) and females *f*_U_, *f*_i_ ((solid lines) as well as infected (green) and uninfected (purple) individuals in a population is shown. In all graphs, we chose a starting frequency of 0.01 for the invading strain, *F*_CI_ =1.3, *T*_CI_ = 0.8 and *β*_CI_ = 0.9. The main difference between the three cases concerns the production of uninfected males, which is decreased by CI in diplodiploids (a), does not change with CI_FM_ (b), and is increased with CI_MD_ (c). Concomitantly, the equilibrium frequency decreases from CI in diplodiploids over CI_FM_ to CI_MD_.

### (3) Turning males into females: similarities and differences between FE and PI

It seems natural to compare FE (in diplodiploids) and PI (in haplodiploids) because in both cases would-be males are transformed into females. Additionally, FE and PI are based on mechanisms that are similar conceptually. PI can be achieved via either a one- or a two-step mechanism (Verhulst *et al*., 2023). In the two-step mechanism, endosymbiont-induced diploidization and feminization are two separate steps (Ma *et al*., 2015), whereas in the one-step mechanism, they co-occur in a manner that cannot be separated (Chen *et al*., 2022; Wang & Verhulst, 2022). The two-step mechanism seems to be necessary whenever diploidization does not automatically lead to feminization, and here, the feminization of diploid males is conceptually identical to the essential role that it plays in FE.

That some manipulations lead to identical infection dynamics is an interesting result of the variability and diversity of the biological mechanisms that reproductive manipulators apply. Given that PI and FE share the same logic, where the parasite benefits if it turns infected males into infected females that offer vertical transmission routes to the endosymbiont, the infection dynamics (equations 6.1-6.4 and 9.1-9.4), invasion conditions (equations 7 & 10) and equilibrium frequencies (equations 8.1-4 & 11.1-4) are identical (Figure 3b, e).

The main conceptual difference between FE and PI is whether fertilization is necessary to produce offspring in the first place. Our equations include this difference: the “successful PI” term of equation 9.1 captures the fact that a proportion of PI infected females can be formed through PI manipulation prior to potential fertilization, whereas FE happens only after fertilization of the eggs (equation 6.1) (see Figures S1, S2). As our results show, this does not induce any differences in the dynamics, except when invading into a population infected with CI, where PI can avoid some of the detrimental effects of mating with CI infected males and invade more easily than FE (compare FE invading into CI in Table 3 with PI invading into CI_FM_ or CI_MD_ in Table 4).

In a natural population a successful feminiser may inadvertently hinder its own transmission by causing a shortage of males and even increase host population extinction risk (Fisher *et al*., 2024). Since the details of sperm limitation depend on male mating capacity (Svensson *et al*., 1998; Wedell *et al*., 2002) and other ecological factors such as the efficiency of mate-finding (deRivera & Vehrencamp, 2001), we did not include them in our minimal model, but only highlight here that transmission of PI endosymbionts does not depend on host matedness, and male success plays no role in the dynamics.

## VII. Useful deviations from the assumptions in the minimal model

While our minimal model has the advantage of making results from different systems directly comparable, it omits real-life factors such as variability in compatibility between different strains, co-infections at the host individual level, and population structure. These deviations extend our minimal framework and aid in explaining the infection frequencies that have been observed in nature, including the coexistence of multiple symbiont strains in one host population. In the following, we first discuss insights from existing model extensions and then consider other factors that can shape natural endosymbiont dynamics as prospects for future model extensions.

### (1) Varying compatibility between different CI strains

We have assumed throughout that crosses between males and females infected with different CI strains are incompatible both ways. With such bidirectional CI, one strain is predicted to outcompete the other, depending on the parameter values and on the initial frequencies (Rousset *et al*., 1991).

However, CI can also be unidirectional: strain A can rescue CI of strain B (leading to the production of viable progeny), but not vice versa (Hoffmann & Turelli, 1988; Kambhampati *et al*., 1993). Hence, strain A is more likely to persist than strain B.

Some degree of compatibility can lead to interesting outcomes. Hurst *et al*. (2002) consider a resident CI symbiont facing invasion of a mutant that has maintained the ability to rescue CI and gained the ability to induce a sex-ratio distorting manipulation (PI, FE or MK). They showed CI to be highly vulnerable to invasion of compatible mutants (Hurst *et al*., 2002), as the compatibility eliminates the priority effect which favoured the resident CI strain. Incompatible invaders experience a different fate: they fail, as they suffer dramatic fecundity losses when mating in a population with high CI prevalence.

### (2) Co-infections at the individual level

A single host individual can be simultaneously infected with multiple strains of endosymbiotic bacteria, which allows a variety of endosymbiotic interactions that affect host biology (Curry *et al*., 2015; White *et al*., 2011). Theory shows that co-infection at the individual level enables coexistence of different reproductive manipulators in a population (Engelstädter *et al*., 2004; Frank, 1998; Zug & Hammerstein, 2018), not only by allowing different strains to persist inside the same individual, but also through imperfect transmission efficiency, which creates singly infected offspring by doubly infected mothers (Zug & Hammerstein 2018).

Co-infection models of reproductive manipulators typically assume that both manipulations occur. The basic rate of increase of both strains then contributes multiplicatively to invasion success from rarity (Engelstädter *et al*., 2004; Zug & Hammerstein, 2018). This combined effect can facilitate invasion of co-infections, as opposed to scenarios without co-infected individuals. Co-infection with a strong MK parasite, for example, will lower the invasion threshold for a CI strain (Engelstädter *et al*., 2004), and a similar effect occurs when CI co-occurs with SD (Wybouw *et al*. 2023, though they did not explicitly model co-infection, but an endosymbiont that is capable of both types of manipulation). In a natural example, a FE and CI co-infection spread rapidly through the CI-infected population (Miyata *et al*., 2024), probably due to compatibility with the local CI-strain.

### (3) Population structure

Population structure significantly changes the predictions from panmictic models. For one, spatial structure can explain the otherwise mysterious high-frequency persistence of endosymbionts that dramatically reduce the number of males (such as male-killers), even in cases where thelytoky is not possible and males are thus strictly necessary for host reproduction. If subpopulations are connected through moderate dispersal, different subpopulations experience sperm limitation in an asynchronous manner, and can be rescued by recurrent immigration of males (Bonte *et al*., 2008; Groenenboom & Hogeweg, 2002). A similar argument can be made for FE and SD, although we know of no spatially structured models involving these manipulation types.

Importantly, complex host population structure can change the outcomes of models for endosymbiont persistence and speed of spread (for CI symbionts, see the review by Engelstädter & Telschow, 2009). For example, drift can help CI overcome the invasion threshold locally, from where it can spread elsewhere (Reuter *et al*., 2008).

Spatial structure also enables stable coexistence of different symbiont strains. Even a small number of subpopulations connected by dispersal aids the coexistence of different CI-strains in the global population, as long as migration rates between subpopulations are sufficiently low (Flor *et al*., 2007; Keeling *et al*., 2003; Telschow *et al*., 2005). Similar results exist for MK, because spatial structure reduces direct competition between strains (Groenenboom & Hogeweg, 2002).

Aside from spatial population structure, models have investigated the impact of age structure and overlapping generations on CI invasion (Rasgon & Scott, 2004). While the age structured population model yields CI equilibrium frequencies that are comparable to models without such structure (Turelli & Hoffmann 1999), the predicted invasion thresholds are higher and depend on the stability of the population age distribution.

### (4) Variability of *T, F* and *β*

Our modelling framework focuses on three components: *F, T* and *β*. It is crucial to remember that their values may vary over time and across different ecological contexts.

Hoffmann *et al*. (1990), for example, observed nearly perfect transmission efficiency *T* for *Wolbachia* infecting *Drosophila simulans* under laboratory conditions, but transmission rates in nature were significantly lower. One possible reason for this difference is the influence of temperature on vertical transmission efficiency (Hague *et al*., 2020; 2022). Whether transmission efficiency increases or decreases with temperature depends on the study system (Martins *et al*., 2023).

Infection-associated fecundity *F* can also be context-dependent. Under low larval density, *Wolbachia* enhances the survivorship of infected female *Aedes albopicture* larvae, while under high density, infected females are less competitive compared to uninfected conspecifics (Gavotte *et al*., 2010). In another example, infected fruit fly females show increased fecundity only under low nutrient conditions (Brownlie *et al*., 2009; Unckless & Jaenike, 2012).

Host defences may impact the efficiency of any reproductive manipulation *β*, and maternal effects can strongly influence the strength of CI (Wybouw *et al*., 2022). If hosts evolve traits that limit or even suppress the reproductive manipulation, the spread of resistant hosts hinders the persistence of the endosymbiont (Charlat *et al*., 2007; Hayashi *et al*., 2018; Koehncke *et al*., 2009; Majerus & Majerus, 2010; Richardson *et al*., 2023). Interestingly, as host suppression of one type of manipulation evolves, some endosymbionts have been shown to switch to another form of manipulation (Hornett *et al*., 2008). Finally, the magnitude of the benefits that a symbiont receives from its manipulation can fluctuate. In the case of MK, for example, the resources freed by the death of MK-infected males are not always redistributed effectively between surviving siblings (Balas *et al*., 1996; Unckless & Jaenicke, 2012), reflecting variation of *Φ*.

Changes in *T, F* or *β* can rapidly impact parasite abundance in a host population. After an initial increase, the prevalence of *Rickettsia* in a *Bemisia tabaci* population rapidly declined, with infected females no longer showing the previously recorded female-biased offspring production and fitness benefits (Himler *et al*., 2011; Bockoven *et al*., 2020). Predicting events such as the loss of endosymbiont manipulation is challenging, and certain interdependencies are unique to different host-parasite interactions, which makes for limited generality. However, when studying a particular host-endosymbiont system, integrating known context-dependent factors into models (example: Fisher et al. 2022) will enhance their predictive power.

### (5) Male fitness effects

Here we highlight the possibility that infected males could experience enhanced fitness, modellable as a parameter equivalent to *F* but expressed in males. The logic behind this is that even though there is no direct selection on maternally transmitted symbionts to increase male fitness, there could be indirect selection to do so. If, for example, male attractiveness was enhanced, then in the paternal-origin cases (i.e., CI) males could impact the reproduction of more females than an assumption of random mating permits them to do, which would benefit the symbiont. Enhanced sperm competitiveness or increased male lifespan would have similar effects.

This reasoning has found theoretical support. Building upon the sperm competition model by Prout & Bundgaard (1977), Hoffmann & Turelli (1997) showed that higher sperm competitiveness of infected males facilitates the spread of CI. However, existing empirical evidence is inconclusive. Studies of the effects of CI-inducing *Wolbachia* on male fertility, including sperm competition, have provided conflicting results, reporting both beneficial (Wade & Chang, 1995; Hariri *et al*., 1998) and detrimental effects (Snook *et al*., 2000; Champion de Crespigny & Wedell, 2006). Likewise, reported effects of CI-inducing *Wolbachia* on male lifespan are inconsistent (Dobson *et al*., 2004; Fry *et al*., 2004; Lewis *et al*., 2011) or negative (Rasgon, 2012). The tradition of building discrete-generation models does not easily allow lifespan modifications to be built into our general framework. Future work could usefully adopt continuous-time lifespan models, which exist for insect reproduction otherwise, e.g. Ekrem & Kokko (2023).

Disassortative mating could similarly have beneficial effects: if CI-infected males targeted uninfected females with courtship, this would benefit the parasite. The question of whether mate preferences have evolved in CI-infected hosts has usually been addressed from the host’s perspective: under CI, there should be selection on hosts to preferentially mate with compatible mates (i.e., infected males with infected females, and uninfected females with uninfected males). While theory suggests that such CI-avoiding mate preferences can indeed evolve (Champion de Crespigny *et al*., 2005), empirical evidence is scarce (Champion de Crespigny & Wedell, 2007; Hoffmann *et al*., 1990; Cruz et al., 2024; but see Vala *et al*., 2004).

The flipside, i.e. the symbiont’s perspective, appears not to have been considered either theoretically on empirically, despite a clear expectation that the parasite benefits if it can counteract assortative mating in the host, making it random or, in the best case (for the parasite), disassortative. We are not aware of this type of parasitic behavioural manipulation existing, even though, in general, manipulation of host behaviour to increase parasite fitness is well documented (Godfrey & Poulin, 2022). There are thus clear empirical as well as theoretical avenues for further work.

## VIII. Contrasting the ease of endosymbiont spread within host populations with their incidence across species

### (1) Predictions about efficiency, prevalence and incidence of different manipulation types

Our general modelling framework allows us to make predictions about the ease with which different types of reproductive manipulation spread through populations. From these, we can extrapolate both their prevalence (proportion of infected individuals in a population) and, with some caveat (see below), their incidence (proportion of infected species). All else being equal, we make three specific predictions.

1. PI can be considered the ‘most efficient’ reproductive manipulation (Verhulst *et al*., 2023; Li *et al*., 2024) because it 1) produces many females, without any fitness costs (in contrast to MK, for example, where males are ‘wasted’), and 2) avoids the problem of creating a shortage of males in the host population (because males are no longer necessary). Based solely on its capacity to spread and remain stable in a host population, even at high prevalence (because the lack of males will not produce any problems), one would expect PI to be the most common reproductive manipulation.
2. Any manipulation that turns what would otherwise become infected dead-end males into infected females (PI, FE, SD) is more efficient than a manipulation that kills infected males. When, for example, comparing FE and MK, initial FE spread only requires some increase in *β*_FE_, while the benefit of MK not only depends on *β*_MK_, but also on the resources transferred from killed brothers to their sisters, *Φ*. Sibling competition (*Φ* > 0) remains a key requirement the success of MK. Not every arthropod species meets this prerequisite, which should constrain the incidence of male-killers. There is no analogous requirement of sibling competition for other known manipulations.
3. Lastly, CI does not facilitate invasion from rarity, which should negatively impact its chances to spread within, and indirectly also between, host populations, compared to other types of reproductive manipulation. However, once established, it can defend itself against invading strains, assuming incompatibility applies in matings between the invader and the resident CI strain; we elaborate on this below.

### (2) The gap between predictions and reality

The expectations outlined above do not seem to match actual incidences of reproductive manipulations in arthropods. CI is considered to be the most common reproductive manipulation (Turelli *et al*., 2022), even though positive frequency dependence makes it difficult for CI to invade when rare (see section IV.5). This suggests that CI-independent fitness benefits (*F*_CI_ > 1) play an important role in overcoming the threshold frequency (Figure 3f; Karisto *et al*., 2022). However, positive frequency dependence also comes with a benefit: CI, once established, is able to resist invasion by other types of (incompatible) manipulators, which highlights that priority effects (section V.2) could play a strong role in explaining CI’s high incidence in nature.

A non-mutually exclusive possibility for the high incidence of CI is based on a clade selection argument: CI-inducing symbionts might be more likely than non-CI-inducing symbionts to spread to new host species because CI-associated positive frequency dependence leads to particularly high equilibrium frequencies, which then increases the likelihood of host switches via interspecific transmission (Hurst & McVean, 1996; Turelli *et al*., 2022). Currently, the relative roles of priority effects and host switches remain unclear.

Additional discrepancies between theoretical predictions and observed incidences of reproductive manipulations can result from differing mechanistic constraints. Some manipulations may be more easily accomplished than others. MK mechanisms, for example, are very diverse across (and even within) host taxa (Harumoto *et al*., 2018; Arai *et al*., 2022; 2023): simply put, any mechanism that kills a male will do for a MK parasite, while transforming males into functional females is a far more specific task. Thus, it is probably easier to realize a high *β*_MK_ than high *β*_FE_ or *β*_PI_. Similarly, PI may be relatively common in haplodiploid hosts because of mechanistic prerequisites that are already in place in haplodiploids and which facilitate parasitic manipulation towards asexual production of females. With haplodiploid sex determination, parthenogenesis already exists (in the form of arrhenotoky), and in many species fertilization is not necessary for egg activation and formation of centrosomes (Engelstädter, 2008). Inducing a switch towards thelytoky is likely to be more challenging (and hence less common) in host species without this prerequisite.

Mechanistic explanations may also boost the clade-level argument for the spread of CI, for it targets conserved mitotic processes (Beckmann *et al*., 2019) that are found in a broad variety of host species. A manipulation mechanism that works in a diverse array of host taxa is clearly expected to facilitate host switches.

### (3) Uncertainties regarding symbiont incidence

It is unclear how widespread the different manipulation types are across different host species. We know of no attempt to list reported incidences of all reproductive manipulations across arthropod species (although there are lists for specific host taxa (Duplouy & Hornett, 2018), endosymbiont species (Werren & O’Neill, 1997) or reproductive manipulations (Kageyama *et al*., 2012; Ma & Schwander, 2017)). Kageyama *et al*. (2012) list more cases of MK than PI or FE, which would contrast with our predictions that MK incidence should be lower than that of PI or FE. However, their list does not claim completeness and the number of recorded cases of reproductive manipulations have increased since then. More recent summaries for e.g. PI include a greater number of cases (Ma & Schwander, 2017), and among haplodiploid arthropod species that had transitioned from sex to parthenogenesis, PI symbionts were found to be causative in 42% of cases (though this is almost certainly an overestimate due to publication bias (van der Kooi *et al*., 2017)). In contrast to all other manipulations, we find only a few documented cases for SD, possibly due to its comparatively recent discovery. Given the great diversity of arthropods (Stork, 2018), the subset of host species that have been screened for endosymbionts and associated reproductive manipulations remains small.

The assumption that prevalence and incidence of reproductive manipulators are positively correlated is not unproblematic because switching to a new host species is fraught with obstacles. Sanaei *et al*. (2021) separate *Wolbachia* host shifts into four different steps: 1) physical transfer to a new host, 2) proliferation in the new host environment, 3) successful maternal transmission, and 4) spread within the new host population. Although the risk of failure lurks in each of these steps, we focus on the last two, as they correspond to the three key components of our framework: vertical transmission efficiency *T* (step 3), manipulation efficiency *β*, and direct fitness effects *F* (both step 4).

Both *T* and *β* are often low in the new host (Sanaei *et al*., 2021), and there are multiple examples where a different reproductive manipulation is expressed in the recipient host upon symbiont transfer from a donor species (reviewed in Hughes & Rasgon, 2014). For example, when a CI-inducing endosymbiont was artificially transferred into a new host species, it caused MK instead, most likely because the mechanisms that induce CI in the original host interact differently with the physiology of the new host (Sasaki *et al*., 2002; Jaenike, 2007). Taken together, there remain important uncertainties regarding the incidence of different manipulation types across arthropods, not least because of incomplete species sampling and the risky nature of host shifts.

## IX. Importance of reproductive manipulations for arthropod evolution

Endosymbionts that manipulate host reproduction are very common in arthropods and seem to persist for prolonged timespans in their host species (Bailly-Bechet *et al*., 2017; Zug *et al*., 2012). The continued presence of reproductive manipulators allows them to be important drivers of arthropod evolution, particularly with respect to host sex determination and sex chromosomes (Cordaux *et al*., 2011). For example, reproductive manipulators have repeatedly triggered the evolution of new sex chromosomes in hosts with female heterogametic (ZW) sex determination. In African Monarch (*Danaus chrysippus*) butterflies, a neo-W chromosome hitchhikes together with a MK *Spiroplasma* (Martin *et al*., 2020), while in the pill bug *Armadillidium vulgare*, a 3 MD insert from a feminizing *Wolbachia* genome functions as a novel female determining sex chromosomal region (Leclercq *et al*., 2016). In the butterfly *Eurema mandarina*, infection with feminizing *Wolbachia* has not only triggered the loss of the W chromosome, but also caused the disruption of Z chromosome inheritance; as a result, *E. mandarina* relies on the symbiont for female development, as removal of the infection results in all-male offspring (Kern *et al*., 2015; Kageyama *et al*., 2017*a*).

The effects of reproductive manipulation on population dynamics can also impact other aspects of host life history, such as mate choice (Randerson *et al*., 2000*a*; Moreau *et al*., 2001; see also section 7.5), evolving infection dependent sperm allocation (Dunn *et al*., 2006) or selecting for a change in dispersal strategies (Goodacre *et al*., 2009). For example, theory predicts that the presence of a MK endosymbiont selects for increased and male biased dispersal patterns in the host (Bonte *et al*., 2008; 2009). Last but not least, sex-ratio distorting symbionts can increase the risk of host population extinction, due to shortage of males (Fisher *et al*., 2024).

## X. Conclusions

1. Several endosymbiont taxa evolved to drastically manipulate the reproduction of their arthropod hosts. Known phenotypes of reproductive manipulation include male-killing (MK), feminization (FE), parthenogenesis induction (PI), sex allocation distortion (SD), and cytoplasmic incompatibility (CI). CI manipulation occurs in offspring of infected fathers (paternal-origin), while all other manipulations stem from the endosymbiont infection in the mother (maternal-origin).
2. Maternal- and paternal-origin effects differ in the theoretical reasoning behind the spread of reproductive manipulators, but in all cases, the relative fecundity of infected females (compared with uninfected ones or those infected with another strain), symbiont transmission efficiency, and efficiency of reproductive manipulation combine to predict the outcome. Predicted infection dynamics are often specific to the type of manipulation but can also be identical across different manipulations (PI & FE) or haplodiploid and diplodiploid hosts species (MK).
3. Theoretical predictions regarding the ease with which reproductive manipulations spread within host populations sometimes contrast with the observed incidence of such manipulations across species. The endosymbiont with the overall higher basic rate of increase is generally expected to outcompete the other strain, but sometimes more realistic assumptions require taking into account varying compatibility, co-infection within a single individual and population structure. Invasion may also occur between manipulation types that do (CI) and do not (other types) involve positive frequency dependence, which complicates the predictions.
4. CI features positive frequency dependence that makes its dynamic differ from others. Its invasion from rare is independent of its efficiency. Positive frequency dependence cause priority effects: while CI invades with difficulty, it enjoys improved stability once established, and it is difficult for incompatible strains to outcompete an established CI strain. This conceivably also impacts the ease of host switching and could contribute to the widespread occurrence of CI in nature.
5. Male-killing is likewise not rare, even though its prerequisite of sibling competition conceivably limits the number of host species in which MK can facilitate its spread. As a whole, discrepancies between theoretical predictions and the abundance of manipulations across species may be more apparent than real, as some manipulations are more easily achieved than others. An additional challenge is evaluating the incidence when taxa, as well as reproductive manipulations, differ in the sampling effort they have received.
6. Endosymbionts that manipulate host reproduction are widespread across arthropods and can remain stable over prolonged periods of time with consequences for arthropod evolution. Future work could unveil the existence of further reproductive manipulations, similar to the recent discovery of SD, whose existence was hypothesized by theoretical work years before. We have also highlighted knowledge gaps, such as fitness effects related to the lifespan or mating success of infected males.

## XI. Acknowledgements

We thank the Swiss National Foundation, the Alexander von Humboldt foundation, and the University of Zürich for funding.

