## Supplementary Material for "Infection dynamics of endosymbionts that manipulate arthropod reproduction"

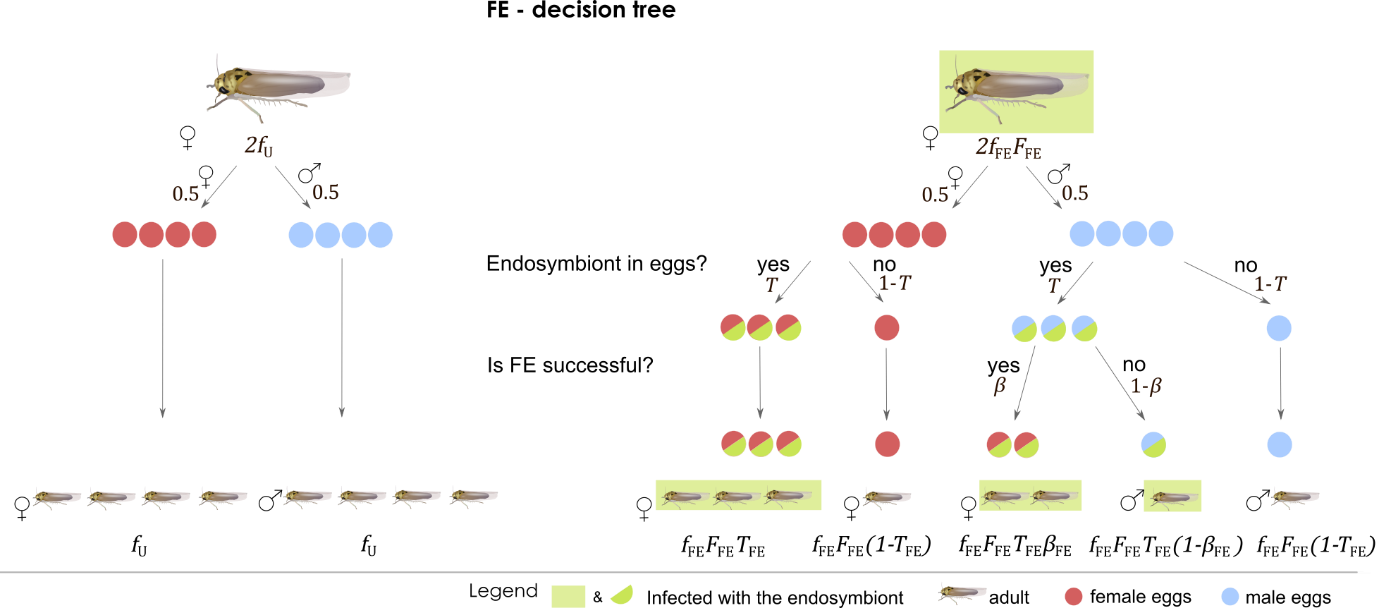

**Figure S1** *Decision tree for feminization (*FE*).* Illustration of offspring proportions for both uninfected (left) and infected (right) females. Green circles and squares indicate infection with the endosymbiont. Blue circles are male eggs; red circles are female eggs.

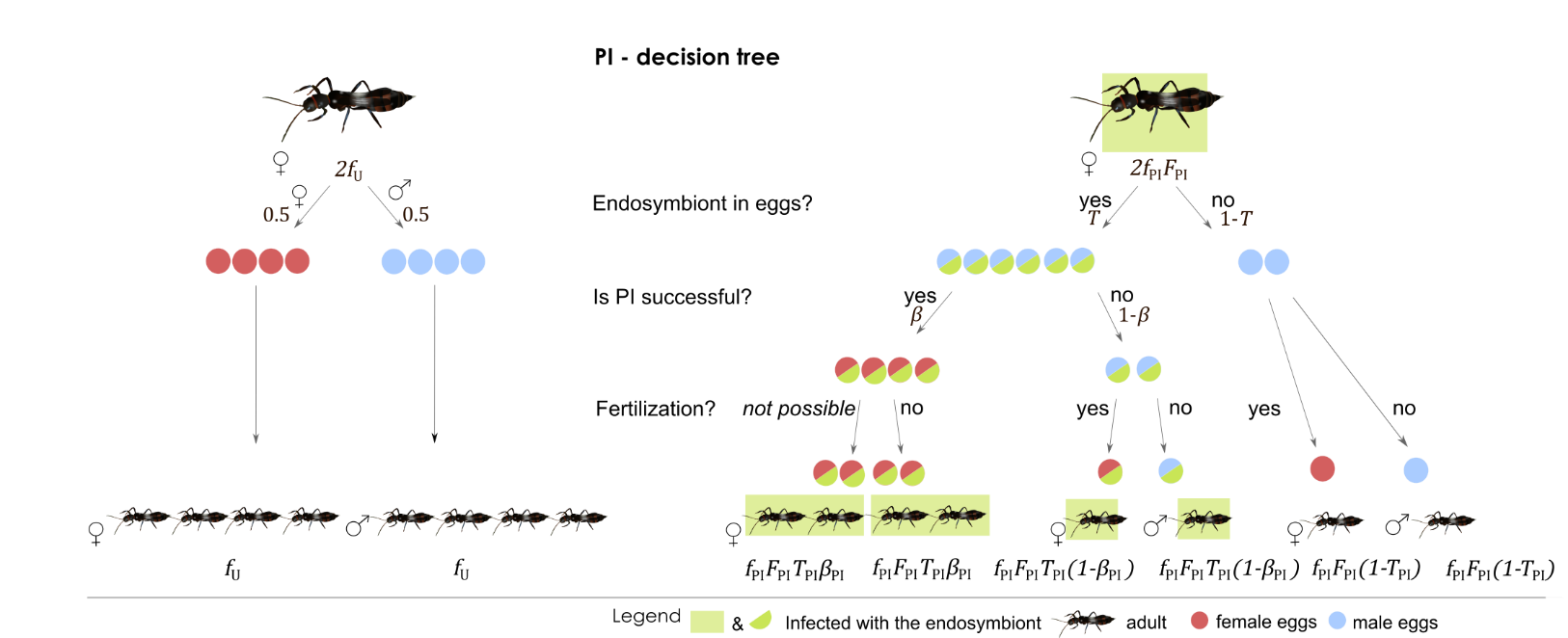

**Figure S2** *Decision tree for parthenogenesis induction (*PI*).* Illustration of offspring proportions for both uninfected (left) and infected (right) females. Green circles and squares indicate infection with the endosymbiont. Blue circles are male eggs; red circles are female eggs. We assume that PI, if successful, takes place immediately after eggs become infected (that is, before potential fertilization), and that fertilization of eggs in which PI was successful is no longer possible. The latter assumption means that all eggs in which PI was successful will develop into infected females.

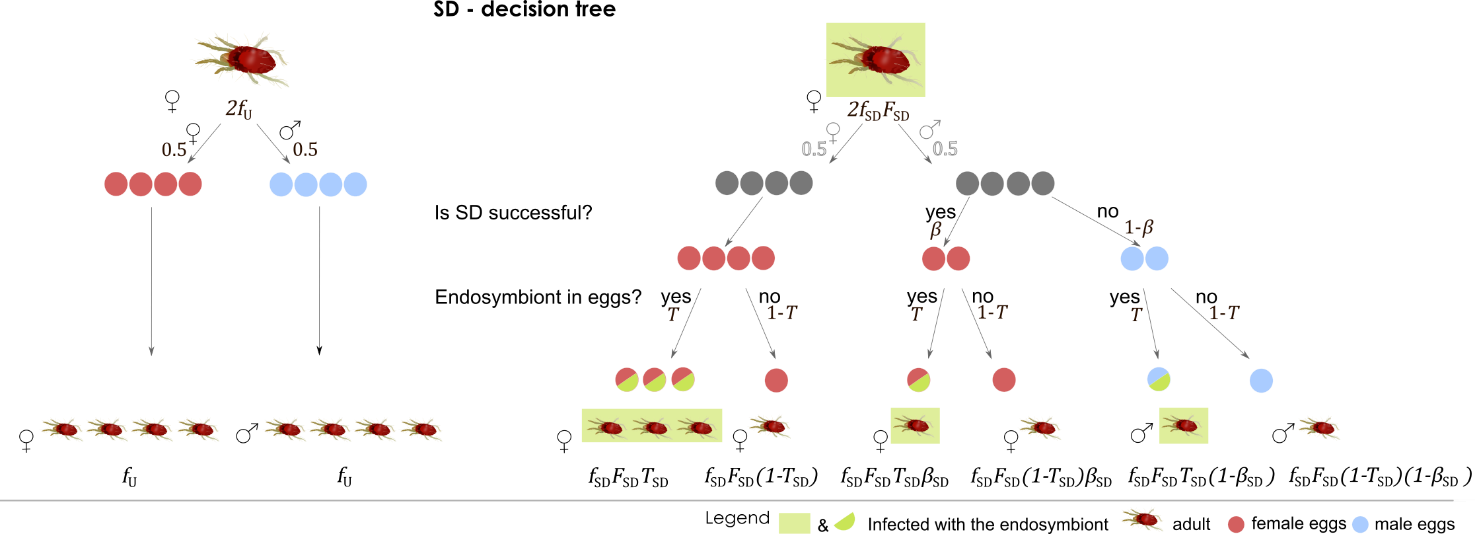

**Figure S3** *Decision tree for sex allocation distortion (*SD*).* Illustration of offspring proportions for both uninfected (left) and infected (right) females. Green circles and squares indicate infection with the endosymbiont. Blue circles are male eggs; red circles are female eggs. Offspring sex ratio depends on the infection status of the mother, not of the offspring; hence SD can take place also in offspring that did not inherit the infection.

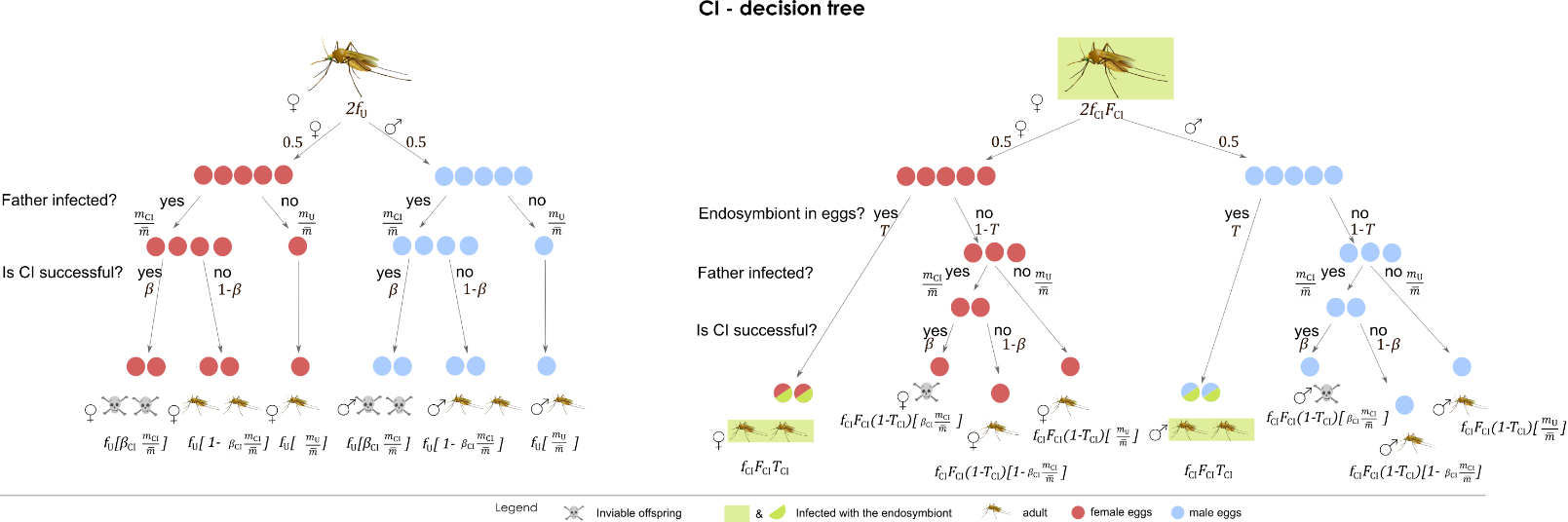

**Figure S4** *Decision tree for cytoplasmic incompatibility (*CI*) in diplodiploid hosts.* Illustration of offspring proportions for both uninfected (left) and infected (right) females. Green circles and squares indicate infection with the endosymbiont. Blue circles are male eggs; red circles are female eggs. A proportion $\beta_{\mathrm{CI}}$ of offspring sired by infected fathers (whose proportion is $\frac{m_{\mathrm{CI}}}{\bar{m}}$, with ${{\bar{m}=m}_{U}+m}_{\mathrm{CI}}$) suffer from CI and die, symbolized by a skull.

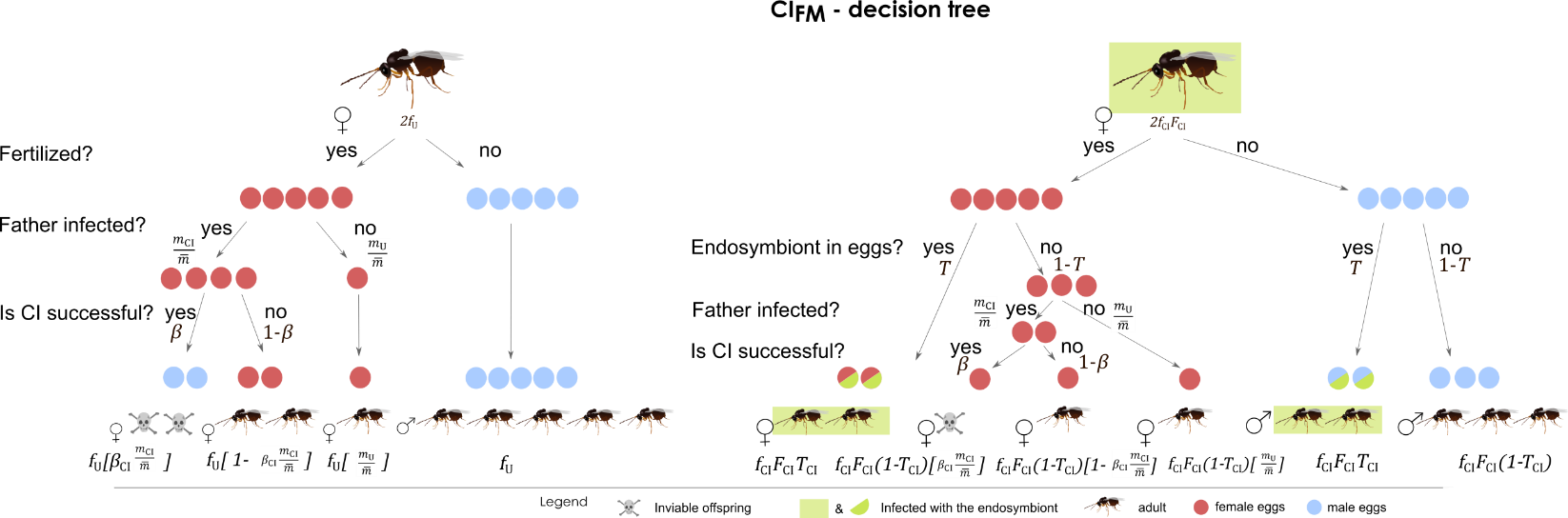

**Figure S5** *Decision tree for cytoplasmic incompatibility with female mortality (*CI_FM_*) in haplodiploid hosts.* Illustration of offspring proportions for both uninfected (left) and infected (right) females. Green circles and squares indicate infection with the endosymbiont. Blue circles are male eggs; red circles are female eggs. A proportion $\beta_{\mathrm{CI}}$ of female progeny sired by infected fathers (whose proportion is $\frac{m_{\mathrm{CI}}}{\bar{m}}$ , with ${{\bar{m}=m}_{U}+m}_{\mathrm{CI}}$) suffer from CI and die, symbolized by a skull. Since uninfected males are produced without fertilization, they do not suffer any CI effects.

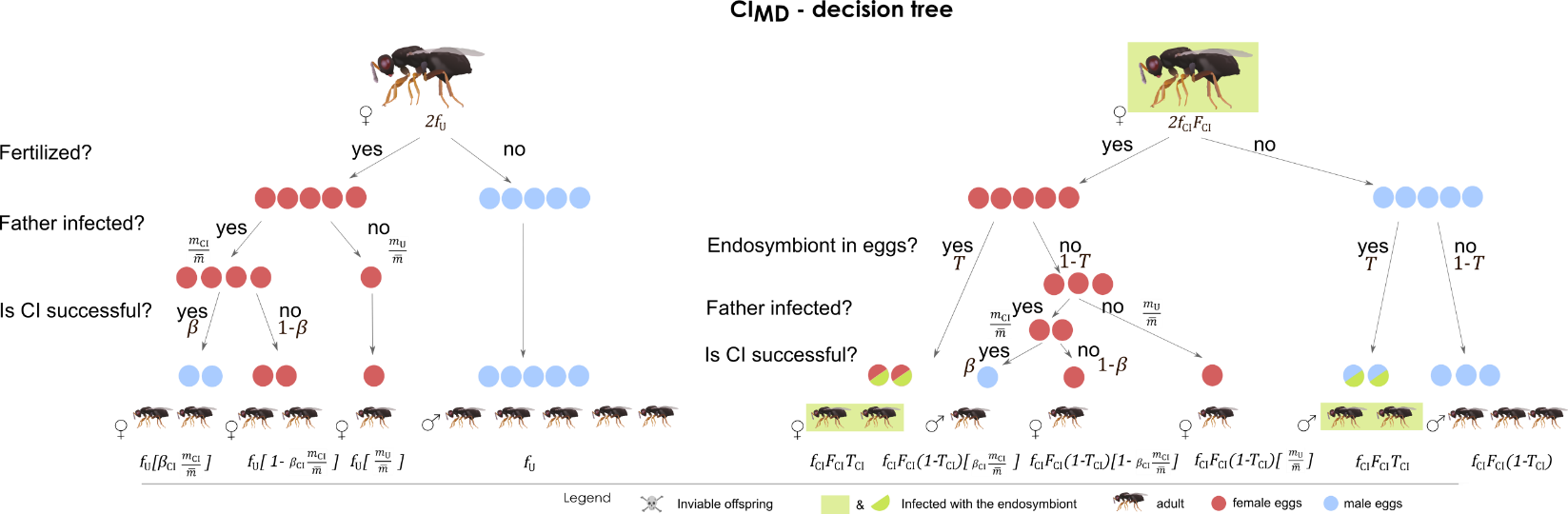

**Figure S6** *Decision tree for cytoplasmic incompatibility with male development (*CI_MD_*) in haplodiploid hosts.* Illustration of offspring proportions for both uninfected (left) and infected (right) females. Green circles and squares indicate infection with the endosymbiont. Blue circles are male eggs; red circles are female eggs. A proportion $\beta_{\mathrm{CI}}$ of female progeny sired by infected fathers (whose proportion is $\frac{m_{\mathrm{CI}}}{\bar{m}}$ with ${{\bar{m}=m}_{U}+m}_{\mathrm{CI}}$) suffer from CI and develop as males. Since uninfected males are produced without fertilization, they do not suffer any CI effects.

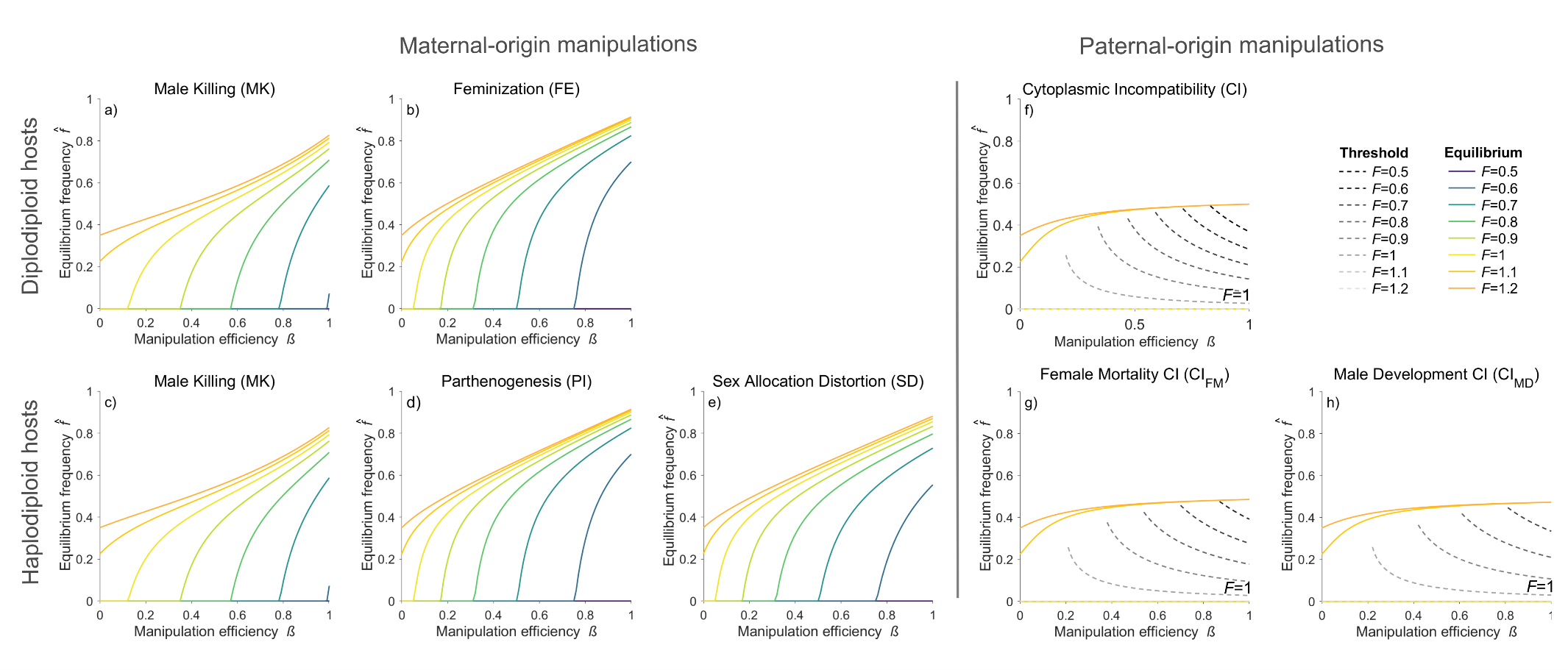
**Figure S7** *Equilibrium frequencies (infected females in the population)* $\hat{f}$(coloured lines) and CI invasion thresholds $\hat{f}_{\mathrm{CI}}^{\mathrm{THR}}$(grey dashed lines) for the different manipulation types, for varying manipulation efficiency *β* (x axis) and relative fecundity *F* (colour). Examples were derived with $T=0.95$ , and, for MK, $\Phi=0.85$. a), b) and f) show results for diplodiploid hosts, and c), d), e), g) and h) for haplodiploid hosts. For all parameter combinations that do not fulfil the CI invasion condition (equations 16, 19 and 22), invasion is not possible and hence the equilibrium frequency is zero, which in these examples applies to all cases where *F* <= 1 (f, g, h). For these cases, the corresponding threshold frequencies are shown (dashed lines). In contrast, for cases where *F* > 1, there is no invasion threshold, i.e., CI symbionts can invade from rarity and will reach the corresponding equilibrium frequency.

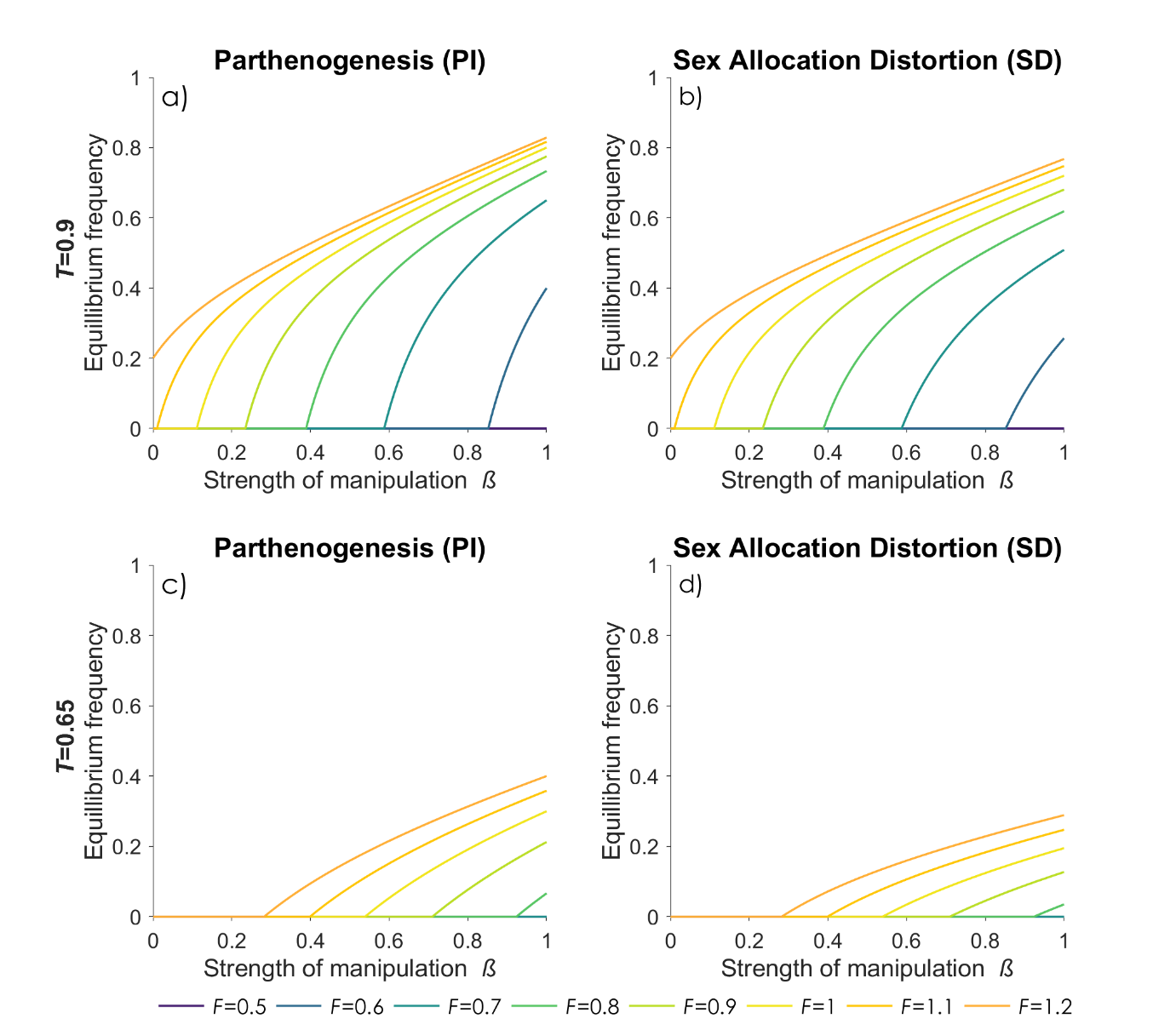

**Figure S8** *Comparison between infected female equilibrium frequencies* $\hat{f}_{i}$*of PI (a, c) and SD (b, d) endosymbionts for different transmission efficiencies (T = 0.9 (a, b), T = 0.65 (c, d)).* Different colours represent different values for relative fecundity *F*. Since invasion conditions for PI and SD are identical, equilibrium frequencies of PI and SD become positive at identical beta values). However, the equilibria then follow different trajectories, with SD reaching lower frequencies for the same parameter combinations. This is a result of SD increasing not only the number of infected, but also uninfected females produced each generation.

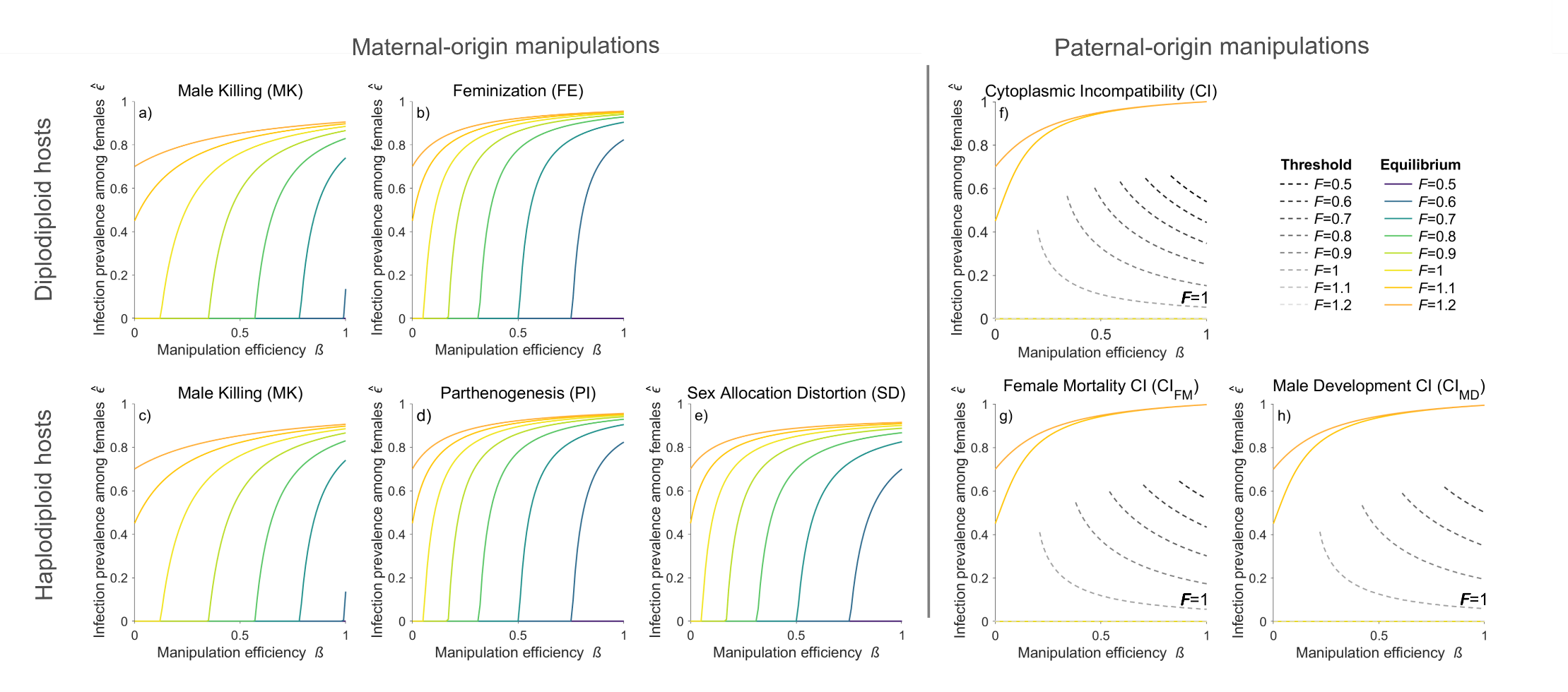

**Figure S9** Equilibrium frequencies (infected females among females) $\hat{\epsilon}$(coloured lines) and CI invasion thresholds $\hat{\varepsilon}_{\mathrm{CI}}^{\mathrm{THR}}$(grey dashed lines) for the different manipulation types, for varying manipulation efficiency *β* (x axis) and relative fecundity *F* (colour). Examples were derived with $T=0.95$ , and, for MK, $\Phi=0.85$. a), b) and f) show results for diplodiploid hosts, and c), d), e), g) and h) for haplodiploid hosts. For all parameter combinations that do not fulfil the CI invasion condition (equations 16, 19 and 22), invasion is not possible and hence the equilibrium frequency is zero, which in these examples applies to all cases where *F* <= 1 (f, g, h). For these cases, the corresponding threshold frequencies are shown (dashed lines). In contrast, for cases where *F* > 1, there is no invasion threshold, i.e., CI symbionts can invade from rarity and will reach the corresponding equilibrium frequency.

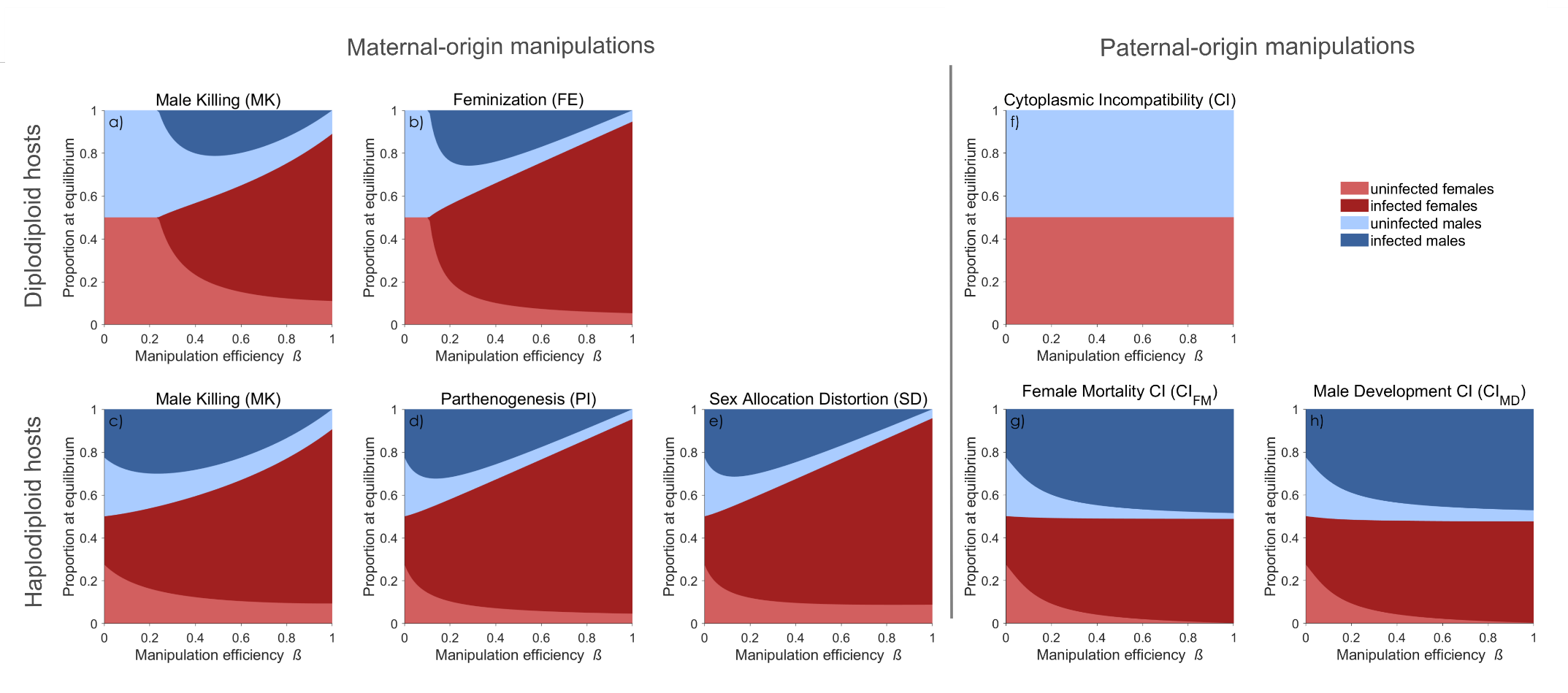

**Figure S10** *Proportion at equilibrium of all four types of individuals in the population* (infected females, $\hat{f}_{i}$; uninfected females, $\hat{f}_{U}$; infected males, $\hat{m}_{i}$; uninfected males, $\hat{m}_{U}$) for the different manipulation types, for varying manipulation efficiency *β* (x axis). Examples were derived with $T=0.95$ , $F=0.95$ and, for MK, $\Phi=0.85$. a), b) and f) show results for diplodiploid hosts, and c), d), e), g) and h) for haplodiploid hosts.

***Supplementary Information Table S1:*** *Diplodiploids - Recursion Equations*

|  | MK | CI | FE |
| --- | --- | --- | --- |
| U | ${f'}_{\mathrm{MK}}=\frac{f_{\mathrm{MK}}F_{\mathrm{MK}}T_{\mathrm{MK}}R}{\overline{\omega}}$  ${f'}_{U}=\frac{f_{U}+f_{\mathrm{MK}}F_{\mathrm{MK}}\left( 1-T_{\mathrm{MK}} \right)R}{\overline{\omega}}$ | ${f'}_{\mathrm{CI}}=\frac{f_{\mathrm{CI}}F_{\mathrm{CI}}T_{\mathrm{CI}}}{\overline{\omega}}$  ${f'}_{U}=\frac{\left( {f_{U}+f}_{\mathrm{CI}}F_{\mathrm{CI}}\left( 1-T_{\mathrm{CI}} \right) \right)\left[ 1-\frac{m_{\mathrm{CI}}\beta_{\mathrm{CI}}}{\overline{m}} \right]}{\overline{\omega}}$ | ${f'}_{\mathrm{FE}}=\frac{f_{\mathrm{FE}}F_{\mathrm{FE}}T_{\mathrm{FE}}\left( 1+\beta_{\mathrm{FE}} \right)}{\overline{\omega}}$  ${f'}_{U}=\frac{{f_{U}+f}_{\mathrm{FE}}F_{\mathrm{FE}}\left( 1-T_{\mathrm{FE}} \right)}{\overline{\omega}}$ |
| MK | ${f'}_{\mathrm{MK}_{1}}=\frac{f_{\mathrm{MK}_{1}}F_{\mathrm{MK}_{1}}T_{\mathrm{MK}_{1}}R_{\mathrm{MK}_{1}}}{\overline{\omega}}$  ${f'}_{{MK}_{2}}=\frac{f_{\mathrm{MK}_{2}}F_{\mathrm{MK}_{2}}T_{\mathrm{MK}_{2}}R_{\mathrm{MK}_{2}}}{\overline{\omega}}$  ${f'}_{U}=\frac{f_{U}+f_{\mathrm{MK}_{1}}{F_{\mathrm{MK}}}_{1}\left( 1-T_{\mathrm{MK}_{1}} \right)R_{\mathrm{MK}_{1}}+f_{\mathrm{MK}_{2}}{F_{\mathrm{MK}}}_{2}\left( 1-T_{{MK}_{2}} \right)R_{\mathrm{MK}_{2}}}{\overline{\omega}}$ |  |  |
| CI | ${f'}_{\mathrm{MK}}=\frac{f_{\mathrm{MK}}F_{\mathrm{MK}}RT_{\mathrm{MK}}\left[ 1-\frac{m_{\mathrm{CI}}\beta_{\mathrm{CI}}}{\overline{m}} \right]}{\overline{\omega}}$  ${f'}_{\mathrm{CI}}=\frac{f_{\mathrm{CI}}F_{\mathrm{CI}}T_{\mathrm{CI}}}{\overline{\omega}}$  ${f'}_{U}=\frac{\left( {f_{U}+f}_{\mathrm{CI}}F_{\mathrm{CI}}\left( 1-T_{\mathrm{CI}} \right)+f_{\mathrm{MK}}F_{\mathrm{MK}}R\left( 1-T_{\mathrm{MK}} \right) \right)\left[ 1-\frac{m_{\mathrm{CI}}\beta_{\mathrm{CI}}}{\overline{m}} \right]}{\overline{\omega}}$ | ${f'}_{\mathrm{CI}_{1}}=\frac{f_{\mathrm{CI}_{1}}F_{\mathrm{CI}_{1}}t_{\mathrm{CI}_{1}}\left[ 1-\frac{m_{\mathrm{CI}_{2}}\beta_{\mathrm{CI}_{2}}}{\overline{m}} \right]}{\overline{\omega}}$  ${f'}_{{CI}_{2}}=\frac{f_{\mathrm{CI}_{2}}F_{\mathrm{CI}_{2}}T_{\mathrm{CI}_{2}}\left[ 1-\frac{m_{\mathrm{CI}_{1}}\beta_{\mathrm{CI}_{1}}}{\overline{m}} \right]}{\overline{\omega}}$  ${f'}_{U}=\frac{\left( {f_{U}+f}_{\mathrm{CI}_{1}}F_{\mathrm{CI}_{1}}\left( 1-T_{\mathrm{CI}_{1}} \right)+f_{\mathrm{CI}_{2}}F_{\mathrm{CI}_{2}}\left( 1-T_{\mathrm{CI}_{2}} \right) \right)\left[ 1-\frac{m_{\mathrm{CI}_{1}}\beta_{\mathrm{CI}_{1}}+m_{\mathrm{CI}_{2}}\beta_{\mathrm{CI}_{2}}}{\overline{m}} \right]}{\overline{\omega}}$ |  |
| FE | ${f'}_{MK}=\frac{f_{\mathrm{MK}}F_{\mathrm{MK}}T_{\mathrm{MK}}R}{\overline{\omega}}$  ${f'}_{\mathrm{FE}}=\frac{f_{\mathrm{FE}}F_{\mathrm{FE}}T_{\mathrm{FE}}\left( 1+\beta_{\mathrm{FE}} \right)}{\overline{\omega}}$  ${f'}_{U}=\frac{f_{U}+f_{\mathrm{MK}}F_{\mathrm{MK}}\left( 1-T_{\mathrm{MK}} \right)R{+f}_{\mathrm{FE}}F_{\mathrm{FE}}\left( 1-T_{\mathrm{FE}} \right)}{\overline{\omega}}$ | ${f'}_{\mathrm{CI}}=\frac{f_{\mathrm{CI}}F_{\mathrm{CI}}T_{\mathrm{CI}}}{\overline{\omega}}$  ${f'}_{\mathrm{FE}}=\frac{f_{\mathrm{FE}}F_{\mathrm{FE}}T_{\mathrm{FE}}\left( 1+\beta_{\mathrm{FE}} \right)\left[ 1-\frac{m_{\mathrm{CI}}\beta_{\mathrm{CI}}}{\overline{m}} \right]}{\overline{\omega}}$  ${f'}_{U}=\frac{\left( {f_{U}+f}_{\mathrm{CI}}F_{\mathrm{CI}}\left( 1-T_{\mathrm{CI}} \right)+f_{\mathrm{FE}}F_{\mathrm{FE}}\left( 1-T_{\mathrm{FE}} \right) \right)\left[ 1-\frac{m_{\mathrm{CI}}\beta_{\mathrm{CI}}}{\overline{m}} \right]}{\overline{\omega}}$ | ${f'}_{\mathrm{FE}_{1}}=\frac{f_{\mathrm{FE}_{1}}F_{\mathrm{FE}_{1}}T_{\mathrm{FE}_{1}}\left( 1+\beta_{\mathrm{FE}_{1}} \right)}{\overline{\omega}}$  ${f'}_{\mathrm{FE}_{2}}=\frac{f_{\mathrm{FE}_{2}}F_{\mathrm{FE}_{2}}T_{\mathrm{FE}_{2}}\left( 1+\beta_{\mathrm{FE}_{2}} \right)}{\overline{\omega}}$  ${f'}_{U}=\frac{f_{U}{+f}_{\mathrm{FE}_{1}}F_{\mathrm{FE}_{1}}\left( 1-T_{\mathrm{FE}_{1}} \right){+f}_{\mathrm{FE}_{2}}F_{\mathrm{FE}_{2}}\left( 1-T_{\mathrm{FE}_{2}} \right)}{\overline{\omega}}$ |

U=uninfected

***Supplementary Information Table S2:*** *Haplodiploids - Recursion Equations*

|  | CI_FM_ |
| --- | --- |
| U | ${f'}_{\mathrm{CI}_{\mathrm{FM}}}=\frac{f_{\mathrm{CI}_{\mathrm{FM}}}F_{\mathrm{CI}_{\mathrm{FM}}}T_{\mathrm{CI}_{\mathrm{FM}}}}{\overline{\omega}}$  ${m'}_{\mathrm{CI}}=\frac{f_{\mathrm{CI}_{\mathrm{FM}}}F_{\mathrm{CI}_{\mathrm{FM}}}T_{\mathrm{CI}_{\mathrm{FM}}}}{\overline{\omega}}$  ${f'}_{U}=\frac{(f_{\mathrm{CI}_{\mathrm{FM}}}F_{\mathrm{CI}_{\mathrm{FM}}}\left( 1-T_{\mathrm{CI}_{\mathrm{FM}}} \right)+f_{U})\left[ 1-\frac{m_{\mathrm{CI}_{\mathrm{FM}}}\beta_{\mathrm{CI}_{\mathrm{FM}}}}{\overline{m}} \right]}{\overline{\omega}}$  ${m'}_{U}=\frac{f_{\mathrm{CI}_{\mathrm{FM}}}F_{\mathrm{CI}_{\mathrm{FM}}}\left( 1-T_{\mathrm{CI}_{\mathrm{FM}}} \right)+f_{U}}{\overline{\omega}}$ |
| CI_FM_ | ${f'}_{\mathrm{CI}_{\mathrm{FM}_{1}}}=\frac{{(f}_{\mathrm{CI}_{\mathrm{FM}_{1}}}F_{\mathrm{CI}_{\mathrm{FM}_{1}}}T_{\mathrm{CI}_{\mathrm{FM}_{1}}})[1-\frac{m_{\mathrm{CI}_{\mathrm{FM}_{2}}}\beta_{\mathrm{CI}_{\mathrm{FM}_{2}}}}{\bar{m}}]}{\bar{\omega}}$  ${m'}_{\mathrm{CI}_{\mathrm{FM}_{1}}}=\frac{f_{\mathrm{CI}_{\mathrm{FM}_{1}}}F_{\mathrm{CI}_{\mathrm{FM}_{1}}}T_{\mathrm{CI}_{\mathrm{FM}_{1}}}}{\bar{\omega}}$  ${f'}_{\mathrm{CI}_{\mathrm{FM}_{2}}}=\frac{{(f}_{\mathrm{CI}_{\mathrm{FM}_{2}}}F_{\mathrm{CI}_{\mathrm{FM}_{2}}}T_{\mathrm{CI}_{\mathrm{FM}_{2}}})[1-\frac{m_{\mathrm{CI}_{\mathrm{FM}_{1}}}\beta_{\mathrm{CI}_{\mathrm{FM}_{1}}}}{\bar{m}}]}{\bar{\omega}}$  ${m'}_{\mathrm{CI}_{\mathrm{FM}_{2}}}=\frac{f_{\mathrm{CI}_{\mathrm{FM}_{2}}}F_{\mathrm{CI}_{\mathrm{FM}_{2}}}T_{\mathrm{CI}_{\mathrm{FM}_{2}}}}{\bar{\omega}}$  ${f'}_{U}=\frac{{(f}_{\mathrm{CI}_{\mathrm{FM}_{1}}}F_{\mathrm{CI}_{\mathrm{FM}_{1}}}\left( 1-T_{\mathrm{CI}_{\mathrm{FM}_{1}}} \right)+{(f}_{\mathrm{CI}_{\mathrm{FM}_{2}}}F_{\mathrm{CI}_{\mathrm{FM}_{2}}}\left( 1-T_{\mathrm{CI}_{\mathrm{FM}_{2}}} \right)+f_{U})[1-\frac{m_{\mathrm{CI}_{\mathrm{FM}_{1}}}\beta_{\mathrm{CI}_{\mathrm{FM}_{1}}}+m_{\mathrm{CI}_{\mathrm{FM}_{2}}}\beta_{\mathrm{CI}_{\mathrm{FM}_{2}}}}{\bar{m}}]}{\bar{\omega}}$  ${m'}_{U}=\frac{f_{\mathrm{CI}_{\mathrm{FM}_{1}}}F_{\mathrm{CI}_{\mathrm{FM}_{1}}}\left( 1-T_{\mathrm{CI}_{\mathrm{FM}_{1}}} \right)+f_{\mathrm{CI}_{\mathrm{FM}_{2}}}F_{\mathrm{CI}_{\mathrm{FM}_{2}}}\left( 1-T_{\mathrm{CI}_{\mathrm{FM}_{2}}} \right)+f_{U}}{\bar{\omega}}$ |
| CI_MD_ | ${f'}_{\mathrm{CI}_{\mathrm{FM}}}=\frac{(f_{\mathrm{CI}_{\mathrm{FM}}}F_{\mathrm{CI}_{\mathrm{FM}}}T_{\mathrm{CI}_{\mathrm{FM}}})\left[ 1-\frac{\beta_{\mathrm{CI}_{\mathrm{MD}}}m_{\mathrm{CI}_{\mathrm{MD}}}}{\overline{m}} \right]}{\overline{\omega}}$  ${m'}_{\mathrm{CI}_{\mathrm{FM}}}=\frac{f_{\mathrm{CI}_{\mathrm{FM}}}F_{\mathrm{CI}_{\mathrm{FM}}}T_{\mathrm{CI}_{\mathrm{FM}}}+(f_{\mathrm{CI}_{\mathrm{FM}}}F_{\mathrm{CI}_{\mathrm{FM}}}T_{\mathrm{CI}_{\mathrm{FM}}})\left[ \frac{m_{\mathrm{CI}_{\mathrm{MD}}}\beta_{\mathrm{CI}_{\mathrm{MD}}}}{\overline{m}} \right]}{\overline{\omega}}$  ${f^{'}}_{\mathrm{CI}_{\mathrm{MD}}}=\frac{f_{\mathrm{CI}_{\mathrm{MD}}}F_{\mathrm{CI}_{\mathrm{MD}}}T_{\mathrm{CI}_{\mathrm{MD}}}\left[ 1-\frac{\beta_{\mathrm{CI}_{\mathrm{FM}}}m_{\mathrm{CI}_{\mathrm{FM}}}}{\overline{m}} \right]}{\overline{\omega}}$  ${m'}_{\mathrm{CI}_{\mathrm{MD}}}=\frac{f_{\mathrm{CI}_{\mathrm{MD}}}F_{\mathrm{CI}_{\mathrm{MD}}}T_{\mathrm{CI}_{\mathrm{MD}}}}{\overline{\omega}}$  ${f^{'}}_{U}=\frac{{(f}_{\mathrm{CI}_{\mathrm{FM}}}F_{\mathrm{CI}_{\mathrm{FM}}}\left( 1-T_{\mathrm{CI}_{\mathrm{FM}}} \right)+f_{U}+{(f}_{\mathrm{CI}_{\mathrm{MD}}}F_{\mathrm{CI}_{\mathrm{MD}}}\left( 1-T_{\mathrm{CI}_{\mathrm{MD}}} \right)\left[ 1-\frac{{\beta_{\mathrm{CI}_{\mathrm{FM}}}m_{\mathrm{CI}_{\mathrm{FM}}}+\beta}_{\mathrm{CI}_{\mathrm{MD}}}m_{\mathrm{CI}_{\mathrm{MD}}}}{\bar{m}} \right]}{\bar{\omega}}$  ${m'}_{U}=\frac{f_{\mathrm{CI}_{\mathrm{FM}}}F_{\mathrm{CI}_{\mathrm{FM}}}\left( 1-T_{\mathrm{CI}_{\mathrm{FM}}} \right)+f_{\mathrm{CI}_{\mathrm{MD}}}F_{\mathrm{CI}_{\mathrm{MD}}}\left( 1-T_{\mathrm{CI}_{\mathrm{MD}}} \right){+f}_{U}+{(f}_{\mathrm{CI}_{\mathrm{FM}}}F_{\mathrm{CI}_{\mathrm{FM}}}\left( 1-T_{\mathrm{CI}_{\mathrm{FM}}} \right)+f_{U}+{(f}_{\mathrm{CI}_{\mathrm{MD}}}F_{\mathrm{CI}_{\mathrm{MD}}}\left( 1-T_{\mathrm{CI}_{\mathrm{MD}}} \right))[\frac{m_{\mathrm{CI}_{\mathrm{MD}}}\beta_{\mathrm{CI}_{\mathrm{MD}}}}{\bar{m}}]}{\bar{\omega}}$ |
| MK | ${f'}_{\mathrm{MK}}=\frac{f_{\mathrm{MK}}F_{\mathrm{MK}}T_{\mathrm{MK}}R[1-\frac{\beta_{\mathrm{CI}_{\mathrm{FM}}}m_{\mathrm{CI}_{\mathrm{FM}}}}{\bar{m}}]}{\bar{\omega}}$  ${m'}_{\mathrm{MK}}=\frac{f_{\mathrm{MK}}F_{\mathrm{MK}}t_{\mathrm{MK}}R{(1-\beta}_{\mathrm{MK}})}{\bar{\omega}}$  ${f'}_{\mathrm{CI}_{\mathrm{FM}}}=\frac{{(f}_{\mathrm{CI}_{\mathrm{FM}}}F_{\mathrm{CI}_{\mathrm{FM}}}T_{\mathrm{CI}_{\mathrm{FM}}})[1]}{\bar{\omega}}$  ${m'}_{\mathrm{CI}_{\mathrm{FM}}}=\frac{f_{\mathrm{CI}_{\mathrm{FM}}}F_{\mathrm{CI}_{\mathrm{FM}}}T_{\mathrm{CI}_{\mathrm{FM}}}}{\bar{\omega}}$  ${f'}_{U}=\frac{{(f}_{\mathrm{CI}_{\mathrm{FM}}}F_{\mathrm{CI}_{\mathrm{FM}}}\left( 1-T_{\mathrm{CI}_{\mathrm{FM}}} \right)+f_{U}+f_{\mathrm{MK}}F_{\mathrm{MK}}\left( 1-T_{\mathrm{MK}} \right)R)[1-\frac{m_{\mathrm{CI}_{\mathrm{FM}}}\beta_{\mathrm{CI}_{\mathrm{FM}}}}{\bar{m}}]}{\bar{\omega}}$  ${m'}_{U}=\frac{f_{\mathrm{CI}_{\mathrm{FM}}}F_{\mathrm{CI}_{\mathrm{FM}}}\left( 1-t_{\mathrm{CI}_{\mathrm{FM}}} \right)+f_{\mathrm{MK}}F_{\mathrm{MK}}\left( 1-T_{\mathrm{MK}} \right)R+f_{U}}{\bar{\omega}}$ |
| PI | ${f'}_{\mathrm{PI}}=\frac{2f_{\mathrm{PI}}F_{\mathrm{PI}}T_{\mathrm{PI}}\beta_{\mathrm{PI}}+f_{\mathrm{PI}}F_{\mathrm{PI}}T_{\mathrm{PI}}(1-\beta_{\mathrm{PI}})[1-\frac{m_{\mathrm{CI}_{\mathrm{FM}}}\beta_{\mathrm{CI}_{\mathrm{FM}}}}{\bar{m}}]}{\bar{\omega}}$  ${m'}_{\mathrm{PI}}=\frac{f_{\mathrm{PI}}F_{\mathrm{PI}}T_{\mathrm{PI}}(1-\beta_{\mathrm{PI}})}{\bar{\omega}}$  ${f'}_{\mathrm{CI}_{\mathrm{FM}}}=\frac{f_{\mathrm{CI}_{\mathrm{FM}}}F_{\mathrm{CI}_{\mathrm{FM}}}T_{\mathrm{CI}_{\mathrm{FM}}}}{\bar{\omega}}$  ${m'}_{\mathrm{CI}_{\mathrm{FM}}}=\frac{f_{\mathrm{CI}_{\mathrm{FM}}}F_{\mathrm{CI}_{\mathrm{FM}}}T_{\mathrm{CI}_{\mathrm{FM}}}}{\bar{\omega}}$  ${f'}_{U}=\frac{{(f}_{\mathrm{CI}_{\mathrm{FM}}}F_{\mathrm{CI}_{\mathrm{FM}}}\left( 1-T_{\mathrm{CI}_{\mathrm{FM}}} \right)+f_{U}+f_{\mathrm{PI}}F_{\mathrm{PI}}\left( 1-T_{\mathrm{PI}} \right))[1-\frac{m_{\mathrm{CI}_{\mathrm{FM}}}\beta_{\mathrm{CI}_{\mathrm{FM}}}}{\bar{m}}]}{\bar{\omega}}$  ${m'}_{U}=\frac{f_{\mathrm{CI}_{\mathrm{FM}}}F_{\mathrm{CI}_{\mathrm{FM}}}\left( 1-T_{\mathrm{CI}_{\mathrm{FM}}} \right)+f_{\mathrm{PI}}F_{\mathrm{PI}}\left( 1-T_{\mathrm{PI}} \right)+f_{U}}{\bar{\omega}}$ |
| SD | ${f'}_{\mathrm{SD}}=\frac{f_{\mathrm{SD}}F_{\mathrm{SD}}T_{\mathrm{SD}}(1+\beta_{\mathrm{SD}})[1-\frac{m_{\mathrm{CI}_{\mathrm{FM}}}\beta_{\mathrm{CI}_{\mathrm{FM}}}}{\bar{m}}]}{\bar{\omega}}$  ${m'}_{\mathrm{SD}}=\frac{f_{\mathrm{SD}}F_{\mathrm{SD}}T_{\mathrm{SD}}(1-\beta_{\mathrm{SD}})}{\bar{\omega}}$  ${f'}_{\mathrm{CI}_{\mathrm{FM}}}=\frac{f_{\mathrm{CI}_{\mathrm{FM}}}F_{\mathrm{CI}_{\mathrm{FM}}}T_{\mathrm{CI}_{\mathrm{FM}}}}{\bar{\omega}}$  ${m'}_{\mathrm{CI}_{\mathrm{FM}}}=\frac{f_{\mathrm{CI}_{\mathrm{FM}}}F_{\mathrm{CI}_{\mathrm{FM}}}T_{\mathrm{CI}_{\mathrm{FM}}}}{\bar{\omega}}$  ${f'}_{U}=\frac{{(f}_{\mathrm{CI}_{\mathrm{FM}}}F_{\mathrm{CI}_{\mathrm{FM}}}\left( 1-T_{\mathrm{CI}_{\mathrm{FM}}} \right)+f_{\mathrm{SD}}F_{\mathrm{SD}}\left( 1-T_{\mathrm{SD}} \right)\left( 1+\beta_{\mathrm{SD}} \right)+f_{U})[1-\frac{m_{\mathrm{CI}_{\mathrm{FM}}}\beta_{\mathrm{CI}_{\mathrm{FM}}}}{\bar{m}}]}{\bar{\omega}}$  ${m'}_{U}=\frac{f_{\mathrm{CI}_{\mathrm{FM}}}F_{\mathrm{CI}_{\mathrm{FM}}}\left( 1-T_{\mathrm{CI}_{\mathrm{FM}}} \right)+f_{\mathrm{SD}}F_{\mathrm{SD}}\left( 1-T_{\mathrm{SD}} \right)\left( 1-\beta_{\mathrm{SD}} \right)+f_{U}}{\bar{\omega}}$ |
|  | CI_MD_ |
| U | ${f^{'}}_{\mathrm{CI}_{\mathrm{MD}}}=\frac{(f_{\mathrm{CI}_{\mathrm{MD}}}F_{\mathrm{CI}_{\mathrm{MD}}}T_{\mathrm{CI}_{\mathrm{MD}}})}{\overline{\omega}}$  ${m'}_{\mathrm{CI}_{\mathrm{MD}}}=\frac{f_{\mathrm{CI}_{\mathrm{MD}}}F_{\mathrm{CI}_{\mathrm{MD}}}T_{\mathrm{CI}_{\mathrm{MD}}}}{\overline{\omega}}$  ${f'}_{U}=\frac{({f_{\mathrm{CI}_{\mathrm{MD}}}F}_{\mathrm{CI}_{\mathrm{MD}}}\left( 1-T_{\mathrm{CI}_{\mathrm{MD}}} \right)+f_{U})\left[ 1-\frac{m_{\mathrm{CI}_{\mathrm{MD}}}\beta_{\mathrm{CI}_{\mathrm{MD}}}}{\overline{m}} \right]}{\overline{\omega}}$  ${m^{'}}_{U}=\frac{f_{\mathrm{CI}_{\mathrm{MD}}}F_{\mathrm{CI}_{\mathrm{MD}}}\left( 1-T_{\mathrm{CI}_{\mathrm{MD}}} \right)+f_{U}+({f_{\mathrm{CI}_{\mathrm{MD}}}F}_{\mathrm{CI}_{\mathrm{MD}}}\left( 1-T_{\mathrm{CI}_{\mathrm{MD}}} \right)+f_{U})\left[ \frac{m_{\mathrm{CI}_{\mathrm{MD}}}\beta_{\mathrm{CI}_{\mathrm{MD}}}}{\overline{m}} \right]}{\overline{\omega}}$ |
| CI_MD_ | ${f^{'}}_{\mathrm{CI}_{\mathrm{MD}_{1}}}=\frac{f_{\mathrm{CI}_{\mathrm{MD}_{1}}}F_{\mathrm{CI}_{\mathrm{MD}_{1}}}T_{\mathrm{CI}_{\mathrm{MD}_{1}}}[1-\frac{m_{\mathrm{CI}_{\mathrm{MD}_{2}}}\beta_{\mathrm{CI}_{\mathrm{MD}_{2}}}}{\bar{m}}]}{\bar{\omega}}$  ${m'}_{\mathrm{CI}_{\mathrm{MD}_{1}}}=\frac{f_{\mathrm{CI}_{\mathrm{MD}_{1}}}F_{\mathrm{CI}_{\mathrm{MD}_{1}}}T_{\mathrm{CI}_{\mathrm{MD}_{1}}}+f_{\mathrm{CI}_{\mathrm{MD}_{1}}}F_{\mathrm{CI}_{\mathrm{MD}_{1}}}T_{\mathrm{CI}_{\mathrm{MD}_{1}}}[\frac{m_{\mathrm{CI}_{\mathrm{MD}_{2}}}\beta_{\mathrm{CI}_{\mathrm{MD}_{2}}}}{\bar{m}}]}{\bar{\omega}}$  ${f^{'}}_{\mathrm{CI}_{\mathrm{MD}_{2}}}=\frac{f_{\mathrm{CI}_{\mathrm{MD}_{2}}}F_{\mathrm{CI}_{\mathrm{MD}_{2}}}T_{\mathrm{CI}_{\mathrm{MD}_{2}}}[1-\frac{m_{\mathrm{CI}_{\mathrm{MD}_{1}}}\beta_{\mathrm{CI}_{\mathrm{MD}_{1}}}}{\bar{m}}]}{\bar{\omega}}$  ${m'}_{\mathrm{CI}_{\mathrm{MD}_{2}}}=\frac{f_{\mathrm{CI}_{\mathrm{MD}_{2}}}F_{\mathrm{CI}_{\mathrm{MD}_{2}}}T_{\mathrm{CI}_{\mathrm{MD}_{2}}}+f_{\mathrm{CI}_{\mathrm{MD}_{2}}}F_{\mathrm{CI}_{\mathrm{MD}_{2}}}T_{\mathrm{CI}_{\mathrm{MD}_{2}}}[\frac{m_{\mathrm{CI}_{\mathrm{MD}_{1}}}\beta_{\mathrm{CI}_{\mathrm{MD}_{1}}}}{\bar{m}}]}{\bar{\omega}}$  ${f'}_{U}=\frac{{(f}_{\mathrm{CI}_{\mathrm{MD}_{1}}}F_{\mathrm{CI}_{\mathrm{MD}_{1}}}\left( 1-T_{\mathrm{CI}_{\mathrm{MD}_{1}}} \right)+f_{\mathrm{CI}_{\mathrm{MD}_{2}}}F_{\mathrm{CI}_{\mathrm{MD}_{2}}}\left( 1-T_{\mathrm{CI}_{\mathrm{MD}_{2}}} \right)+f_{U})[1-\frac{m_{\mathrm{CI}_{\mathrm{MD}_{1}}}\beta_{\mathrm{CI}_{\mathrm{MD}_{1}}}+m_{\mathrm{CI}_{\mathrm{MD}_{2}}}\beta_{\mathrm{CI}_{\mathrm{MD}_{2}}}}{\bar{m}}]}{\bar{\omega}}$  ${m^{'}}_{U}=\frac{f_{\mathrm{CI}_{\mathrm{MD}_{1}}}F_{\mathrm{CI}_{\mathrm{MD}_{1}}}\left( 1-T_{\mathrm{CI}_{\mathrm{MD}_{1}}} \right)+f_{\mathrm{CI}_{\mathrm{MD}_{2}}}F_{\mathrm{CI}_{\mathrm{MD}_{2}}}\left( 1-T_{\mathrm{CI}_{\mathrm{MD}_{2}}} \right)+f_{U}+{(f}_{\mathrm{CI}_{\mathrm{MD}_{1}}}F_{\mathrm{CI}_{\mathrm{MD}_{1}}}\left( 1-T_{\mathrm{CI}_{\mathrm{MD}_{1}}} \right)+f_{U}+f_{\mathrm{CI}_{\mathrm{MD}_{2}}}F_{\mathrm{CI}_{\mathrm{MD}_{2}}}\left( 1-T_{\mathrm{CI}_{\mathrm{MD}_{2}}} \right))[\frac{m_{\mathrm{CI}_{\mathrm{MD}_{1}}}\beta_{\mathrm{CI}_{\mathrm{MD}_{1}}}+m_{\mathrm{CI}_{\mathrm{MD}_{2}}}\beta_{\mathrm{CI}_{\mathrm{MD}_{2}}}}{\bar{m}}]}{\bar{\omega}}$ |
| MK | ${f'}_{\mathrm{MK}}=\frac{f_{\mathrm{MK}}F_{\mathrm{MK}}T_{\mathrm{MK}}R[1-\frac{m_{\mathrm{CI}_{\mathrm{MD}}}\beta_{\mathrm{CI}_{\mathrm{MD}}}}{\bar{m}}]}{\bar{\omega}}$  ${m'}_{\mathrm{MK}}=\frac{f_{\mathrm{MK}}F_{\mathrm{MK}}T_{\mathrm{MK}}R{(1-\beta}_{\mathrm{MK}})+f_{\mathrm{MK}}F_{\mathrm{MK}}T_{\mathrm{MK}}R[\frac{m_{\mathrm{CI}_{\mathrm{MD}}}\beta_{\mathrm{CI}_{\mathrm{MD}}}}{\bar{m}}]}{\bar{\omega}}$  ${f^{'}}_{\mathrm{CI}_{\mathrm{MD}}}=\frac{f_{\mathrm{CI}_{\mathrm{MD}}}F_{\mathrm{CI}_{\mathrm{MD}}}T_{\mathrm{CI}_{\mathrm{MD}}}}{\bar{\omega}}$  ${m'}_{\mathrm{CI}_{\mathrm{MD}}}=\frac{f_{\mathrm{CI}_{\mathrm{MD}}}F_{\mathrm{CI}_{\mathrm{MD}}}T_{\mathrm{CI}_{\mathrm{MD}}}}{\bar{\omega}}$  ${f'}_{U}=\frac{f_{\mathrm{MK}}F_{\mathrm{MK}}\left( 1-T_{\mathrm{MK}} \right)R+f_{\mathrm{CI}_{\mathrm{MD}}}F_{\mathrm{CI}_{\mathrm{MD}}}\left( 1-T_{\mathrm{CI}_{\mathrm{MD}}} \right)+f_{U}[1-\frac{m_{\mathrm{CI}_{\mathrm{MD}}}\beta_{\mathrm{CI}_{\mathrm{MD}}}}{\bar{m}}]}{\bar{\omega}}$  ${m'}_{U}=\frac{f_{\mathrm{MK}}F_{\mathrm{MK}}\left( 1-T_{\mathrm{MK}} \right)R+f_{\mathrm{CI}_{\mathrm{MD}}}F_{\mathrm{CI}_{\mathrm{MD}}}\left( 1-T_{\mathrm{CI}_{\mathrm{MD}}} \right)+f_{U}+{(f_{\mathrm{MK}}F_{\mathrm{MK}}\left( 1-T_{\mathrm{MK}} \right)R+f}_{\mathrm{CI}_{\mathrm{MD}}}F_{\mathrm{CI}_{\mathrm{MD}}}\left( 1-T_{\mathrm{CI}_{\mathrm{MD}}} \right)+f_{U})[\frac{m_{\mathrm{CI}_{\mathrm{MD}}}\beta_{\mathrm{CI}_{\mathrm{MD}}}}{\bar{m}}]}{\bar{\omega}}$ |
| PI | ${f'}_{\mathrm{PI}}=\frac{2f_{\mathrm{PI}}F_{\mathrm{PI}}T_{\mathrm{PI}}\beta_{\mathrm{PI}}+f_{\mathrm{PI}}F_{\mathrm{PI}}T_{\mathrm{PI}}(1-\beta_{\mathrm{PI}})[1-\frac{m_{\mathrm{CI}_{\mathrm{MD}}}\beta_{\mathrm{CI}_{\mathrm{MD}}}}{\bar{m}}]}{\bar{\omega}}$  ${m'}_{\mathrm{PI}}=\frac{f_{\mathrm{PI}}F_{\mathrm{PI}}T_{\mathrm{PI}}(1-\beta_{\mathrm{PI}})}{\bar{\omega}}$  ${f^{'}}_{\mathrm{CI}_{\mathrm{MD}}}=\frac{f_{\mathrm{CI}_{\mathrm{MD}}}F_{\mathrm{CI}_{\mathrm{MD}}}T_{\mathrm{CI}_{\mathrm{MD}}}}{\bar{\omega}}$  ${m'}_{\mathrm{CI}_{\mathrm{MD}}}=\frac{f_{\mathrm{CI}_{\mathrm{MD}}}F_{\mathrm{CI}_{\mathrm{MD}}}T_{\mathrm{CI}_{\mathrm{MD}}}}{\bar{\omega}}$  ${f'}_{U}=\frac{{(f}_{\mathrm{CI}_{\mathrm{MD}}}F_{\mathrm{CI}_{\mathrm{MD}}}\left( 1-T_{\mathrm{CI}_{\mathrm{MD}}} \right)+f_{U}+f_{\mathrm{PI}}F_{\mathrm{PI}}{(1-T}_{\mathrm{PI}}))[1-\frac{m_{\mathrm{CI}_{\mathrm{MD}}}\beta_{\mathrm{CI}_{\mathrm{MD}}}}{\bar{m}}]}{\bar{\omega}}$  ${m^{'}}_{U}=\frac{f_{\mathrm{CI}_{\mathrm{MD}}}F_{\mathrm{CI}_{\mathrm{MD}}}\left( 1-T_{\mathrm{CI}_{\mathrm{MD}}} \right)+f_{U}+{(f}_{\mathrm{CI}_{\mathrm{MD}}}F_{\mathrm{CI}_{\mathrm{MD}}}\left( 1-T_{\mathrm{CI}_{\mathrm{MD}}} \right)+f_{U}+f_{\mathrm{PI}}F_{\mathrm{PI}}{(1-T}_{\mathrm{PI}}))\left[ \frac{m_{\mathrm{CI}_{\mathrm{MD}}}}{\bar{m}}\beta_{\mathrm{CI}_{\mathrm{MD}}} \right]}{\bar{\omega}}$ |
| SD | ${f'}_{SD}=\frac{f_{\mathrm{SD}}F_{\mathrm{SD}}T_{\mathrm{SD}}(1+\beta_{\mathrm{SD}})[1-\frac{m_{\mathrm{CI}_{\mathrm{MD}}}\beta_{\mathrm{CI}_{\mathrm{MD}}}}{\bar{m}}]}{\bar{\omega}}$  ${m'}_{\mathrm{SD}}=\frac{{2f}_{\mathrm{SD}}F_{\mathrm{SD}}T_{\mathrm{SD}}\left( 1-\beta_{\mathrm{SD}} \right)+(f_{\mathrm{SD}}F_{\mathrm{SD}}T_{\mathrm{SD}}(1+\beta_{\mathrm{SD}})[\frac{m_{\mathrm{CI}_{\mathrm{MD}}}\beta_{\mathrm{CI}_{\mathrm{MD}}}}{\bar{m}}])}{\bar{\omega}}$  ${f^{'}}_{\mathrm{CI}_{\mathrm{MD}}}=\frac{f_{\mathrm{CI}_{\mathrm{MD}}}F_{\mathrm{CI}_{\mathrm{MD}}}T_{\mathrm{CI}_{\mathrm{MD}}}}{\bar{\omega}}$  ${m'}_{\mathrm{CI}_{\mathrm{MD}}}=\frac{f_{\mathrm{CI}_{\mathrm{MD}}}F_{\mathrm{CI}_{\mathrm{MD}}}T_{\mathrm{CI}_{\mathrm{MD}}}}{\bar{\omega}}$  ${f'}_{U}=\frac{{(f}_{\mathrm{CI}_{\mathrm{MD}}}F_{\mathrm{CI}_{\mathrm{MD}}}\left( 1-T_{\mathrm{CI}_{\mathrm{MD}}} \right)+f_{\mathrm{SD}}F_{\mathrm{SD}}\left( 1-T_{\mathrm{SD}} \right){(1+\beta}_{\mathrm{SD}})+f_{U})[1-\frac{m_{\mathrm{CI}_{\mathrm{MD}}}\beta_{\mathrm{CI}_{\mathrm{MD}}}}{\bar{m}}]}{\bar{\omega}}$  ${m^{'}}_{U}=\frac{f_{\mathrm{CI}_{\mathrm{MD}}}F_{\mathrm{CI}_{\mathrm{MD}}}\left( 1-T_{\mathrm{CI}_{\mathrm{MD}}} \right)+f_{\mathrm{SD}}F_{\mathrm{SD}}\left( 1-T_{\mathrm{SD}} \right)\left( 1-\beta_{\mathrm{SD}} \right)+f_{U}+{((f}_{\mathrm{CI}_{\mathrm{MD}}}F_{\mathrm{CI}_{\mathrm{MD}}}\left( 1-T_{\mathrm{CI}_{\mathrm{MD}}} \right)+f_{\mathrm{SD}}F_{\mathrm{SD}}\left( 1-T_{\mathrm{SD}} \right){(1+\beta}_{\mathrm{SD}})+f_{U})[\frac{m_{\mathrm{CI}_{\mathrm{MD}}}\beta_{\mathrm{CI}_{\mathrm{MD}}}}{\bar{m}}])}{\bar{\omega}}$ |
|  | MK |
| U | ${f'}_{\mathrm{MK}}=\frac{f_{\mathrm{MK}}F_{\mathrm{MK}}T_{\mathrm{MK}}R}{\overline{\omega}}$  ${m'}_{\mathrm{MK}}=\frac{f_{\mathrm{MK}}F_{\mathrm{MK}}T_{\mathrm{MK}}R(1-\beta_{\mathrm{MK}})}{\overline{\omega}}$  ${f'}_{U}=\frac{f_{\mathrm{MK}}F_{\mathrm{MK}}\left( 1-T_{\mathrm{MK}} \right)R+f_{U}}{\overline{\omega}}$  ${m'}_{U}=\frac{f_{\mathrm{MK}}F_{\mathrm{MK}}\left( 1-T_{\mathrm{MK}} \right)R+f_{U}}{\overline{\omega}}$ |
| MK | ${f'}_{\mathrm{MK}_{1}}=\frac{f_{\mathrm{MK}_{1}}F_{\mathrm{MK}_{1}}T_{\mathrm{MK}_{1}}R_{\mathrm{MK}_{1}}}{\bar{\omega}}$  ${m'}_{\mathrm{MK}_{1}}=\frac{f_{\mathrm{MK}_{1}}F_{\mathrm{MK}_{1}}T_{\mathrm{MK}_{1}}{R_{\mathrm{MK}_{1}}(1-\beta}_{\mathrm{MK}_{1}})}{\bar{\omega}}$  ${f'}_{\mathrm{MK}_{2}}=\frac{f_{\mathrm{MK}_{2}}F_{\mathrm{MK}_{2}}T_{\mathrm{MK}_{2}}R_{\mathrm{MK}_{2}}}{\bar{\omega}}$  ${m'}_{\mathrm{MK}_{2}}=\frac{f_{\mathrm{MK}_{2}}F_{\mathrm{MK}_{2}}T_{\mathrm{MK}_{2}}{R_{\mathrm{MK}_{2}}(1-\beta}_{\mathrm{MK}_{2}})}{\bar{\omega}}$  ${f'}_{U}=\frac{f_{\mathrm{MK}_{1}}F_{\mathrm{MK}_{1}}\left( 1-T_{\mathrm{MK}_{1}} \right)R_{\mathrm{MK}_{1}}+f_{\mathrm{MK}_{2}}F_{\mathrm{MK}_{2}}\left( 1-T_{\mathrm{MK}_{2}} \right)R_{\mathrm{MK}_{2}}+f_{U}}{\bar{\omega}}$  ${m'}_{U}=\frac{f_{\mathrm{MK}_{1}}F_{\mathrm{MK}_{1}}\left( 1-T_{\mathrm{MK}_{1}} \right)R_{\mathrm{MK}_{1}}+f_{\mathrm{MK}_{2}}F_{\mathrm{MK}_{2}}\left( 1-T_{\mathrm{MK}_{2}} \right)R_{\mathrm{MK}_{2}}+f_{U}}{\bar{\omega}}$ |
| PI | ${f'}_{\mathrm{MK}}=\frac{f_{\mathrm{MK}}F_{\mathrm{MK}}T_{\mathrm{MK}}R}{\bar{\omega}}$  ${m'}_{\mathrm{MK}}=\frac{f_{\mathrm{MK}}F_{\mathrm{MK}}T_{\mathrm{MK}}R{(1-\beta}_{\mathrm{MK}})}{\bar{\omega}}$  ${f'}_{\mathrm{PI}}=\frac{{2f}_{\mathrm{PI}}F_{\mathrm{PI}}T_{\mathrm{PI}}\beta_{\mathrm{PI}}+f_{\mathrm{PI}}F_{\mathrm{PI}}T_{\mathrm{PI}}(1-\beta_{\mathrm{PI}})}{\bar{\omega}}$  ${m'}_{\mathrm{PI}}=\frac{f_{\mathrm{PI}}F_{\mathrm{PI}}T_{\mathrm{PI}}(1-\beta_{\mathrm{PI}})}{\bar{\omega}}$  ${f'}_{U}=\frac{f_{\mathrm{MK}}F_{\mathrm{MK}}\left( 1-T_{\mathrm{MK}} \right)R+f_{\mathrm{PI}}F_{\mathrm{PI}}\left( 1-T_{\mathrm{PI}} \right)+f_{U}}{\bar{\omega}}$  ${m'}_{U}=\frac{f_{\mathrm{MK}}F_{\mathrm{MK}}\left( 1-T_{\mathrm{MK}} \right)R+f_{\mathrm{PI}}F_{\mathrm{PI}}\left( 1-T_{\mathrm{PI}} \right)+f_{U}}{\bar{\omega}}$ |
| SD | ${f'}_{\mathrm{SD}}=\frac{f_{\mathrm{SD}}F_{\mathrm{SD}}T_{\mathrm{SD}}{(1+\beta}_{\mathrm{SD}})}{\bar{\omega}}$  ${m'}_{\mathrm{SD}}=\frac{f_{\mathrm{SD}}F_{\mathrm{SD}}T_{\mathrm{SD}}(1-\beta_{\mathrm{SD}})}{\bar{\omega}}$  ${f'}_{\mathrm{MK}}=\frac{f_{\mathrm{MK}}F_{\mathrm{MK}}T_{\mathrm{MK}}R}{\bar{\omega}}$  ${m'}_{\mathrm{MK}}=\frac{f_{\mathrm{MK}}F_{\mathrm{MK}}T_{\mathrm{MK}}R{(1-\beta}_{\mathrm{MK}})}{\bar{\omega}}$  ${f'}_{U}=\frac{f_{\mathrm{SD}}F_{\mathrm{SD}}\left( 1-T_{\mathrm{SD}} \right)(1+\beta_{\mathrm{SD}})+f_{\mathrm{MK}}F_{\mathrm{MK}}\left( 1-T_{\mathrm{MK}} \right)R+f_{U}}{\bar{\omega}}$  ${m'}_{U}=\frac{f_{\mathrm{SD}}F_{\mathrm{SD}}\left( 1-T_{\mathrm{SD}} \right)\left( 1-\beta_{\mathrm{SD}} \right)+f_{\mathrm{MK}}F_{\mathrm{MK}}\left( 1-T_{\mathrm{MK}} \right)R+f_{U}}{\bar{\omega}}$ |
|  | PI |
| U | ${f'}_{\mathrm{PI}}=\frac{{2f}_{\mathrm{PI}}F_{\mathrm{PI}}T_{\mathrm{PI}}\beta_{\mathrm{PI}}+f_{\mathrm{PI}}F_{\mathrm{PI}}T_{\mathrm{PI}}\left( 1-\beta_{\mathrm{PI}} \right)}{\overline{\omega}}$  ${m'}_{\mathrm{PI}}=\frac{f_{\mathrm{PI}}F_{\mathrm{PI}}T_{\mathrm{PI}}\left( 1-\beta_{\mathrm{PI}} \right)}{\overline{\omega}}$  ${f'}_{U}=\frac{f_{\mathrm{PI}}F_{\mathrm{PI}}\left( 1-T_{\mathrm{PI}} \right)+f_{U}}{\overline{\omega}}$  ${m'}_{U}=\frac{f_{\mathrm{PI}}F_{\mathrm{PI}}\left( 1-T_{\mathrm{PI}} \right)+f_{U}}{\overline{\omega}}$ |
| PI | ${f'}_{\mathrm{PI}_{1}}=\frac{2f_{\mathrm{PI}_{1}}F_{\mathrm{PI}_{1}}T_{\mathrm{PI}_{1}}\beta_{\mathrm{PI}_{1}}+f_{\mathrm{PI}_{1}}F_{\mathrm{PI}_{1}}T_{\mathrm{PI}_{1}}(1-\beta_{\mathrm{PI}_{1}})}{\bar{\omega}}$  ${m'}_{\mathrm{PI}_{1}}=\frac{f_{\mathrm{PI}_{1}}F_{\mathrm{PI}_{1}}T_{\mathrm{PI}_{1}}(1-\beta_{\mathrm{PI}_{1}})}{\bar{\omega}}$  ${f'}_{\mathrm{PI}_{2}}=\frac{{2f}_{\mathrm{PI}_{2}}F_{\mathrm{PI}_{2}}T_{\mathrm{PI}_{2}}\beta_{\mathrm{PI}_{2}}+f_{\mathrm{PI}_{2}}F_{\mathrm{PI}_{2}}T_{\mathrm{PI}_{2}}(1-\beta_{\mathrm{PI}_{2}})}{\bar{\omega}}$  ${m'}_{\mathrm{PI}_{2}}=\frac{f_{\mathrm{PI}_{2}}F_{\mathrm{PI}_{2}}T_{\mathrm{PI}_{2}}(1-\beta_{\mathrm{PI}_{2}})}{\bar{\omega}}$  ${f'}_{U}=\frac{f_{\mathrm{PI}_{1}}F_{\mathrm{PI}_{1}}\left( 1-T_{\mathrm{PI}_{1}} \right)+f_{\mathrm{PI}_{2}}F_{\mathrm{PI}_{2}}\left( 1-T_{\mathrm{PI}_{2}} \right)+f_{U}}{\bar{\omega}}$  ${m'}_{U}=\frac{f_{\mathrm{PI}_{1}}F_{\mathrm{PI}_{1}}\left( 1-T_{\mathrm{PI}_{1}} \right)+f_{\mathrm{PI}_{2}}F_{\mathrm{PI}_{2}}\left( 1-T_{\mathrm{PI}_{2}} \right)+f_{U}}{\bar{\omega}}$ |
| SD | ${f^{'}}_{\mathrm{SD}}=\frac{f_{\mathrm{SD}}F_{\mathrm{SD}}T_{\mathrm{SD}}{(1+\beta}_{\mathrm{SD}})}{\bar{\omega}}$  ${m'}_{\mathrm{SD}}=\frac{f_{\mathrm{SD}}F_{\mathrm{SD}}T_{\mathrm{SD}}(1-\beta_{\mathrm{SD}})}{\bar{\omega}}$  ${f'}_{\mathrm{PI}}=\frac{{2f}_{\mathrm{PI}}F_{\mathrm{PI}}T_{\mathrm{PI}}\beta_{\mathrm{PI}}+f_{\mathrm{PI}}F_{\mathrm{PI}}T_{\mathrm{PI}}(1-\beta_{\mathrm{PI}})}{\bar{\omega}}$  ${m'}_{\mathrm{PI}}=\frac{f_{\mathrm{PI}}F_{\mathrm{PI}}T_{\mathrm{PI}}(1-\beta_{\mathrm{PI}})}{\bar{\omega}}$  ${f'}_{U}=\frac{f_{\mathrm{PI}}F_{\mathrm{PI}}\left( 1-T_{\mathrm{PI}} \right)+f_{\mathrm{SD}}F_{\mathrm{SD}}\left( 1-T_{\mathrm{SD}} \right){(1+\beta}_{\mathrm{SD}})+f_{U}}{\bar{\omega}}$  ${m'}_{U}=\frac{f_{\mathrm{PI}}F_{\mathrm{PI}}\left( 1-T_{\mathrm{PI}} \right)+f_{\mathrm{SD}}F_{\mathrm{SD}}\left( 1-T_{\mathrm{SD}} \right)\left( 1-\beta_{\mathrm{SD}} \right)+f_{U}}{\bar{\omega}}$ |
|  | SD |
| U | ${f'}_{\mathrm{SD}}=\frac{f_{\mathrm{SD}}F_{\mathrm{SD}}T_{\mathrm{SD}}\left( 1+\beta_{\mathrm{SD}} \right)}{\overline{\omega}}$  ${m'}_{\mathrm{SD}}=\frac{f_{\mathrm{SD}}F_{\mathrm{SD}}T_{\mathrm{SD}}\left( 1-\beta_{\mathrm{SD}} \right)}{\overline{\omega}}$  ${f'}_{U}=\frac{f_{\mathrm{SD}}F_{\mathrm{SD}}\left( 1-T_{\mathrm{SD}} \right)\left( 1+\beta_{\mathrm{SD}} \right)+f_{U}}{\overline{\omega}}$  ${m'}_{U}=\frac{f_{\mathrm{SD}}F_{\mathrm{SD}}\left( 1-T_{\mathrm{SD}} \right)\left( 1-\beta_{\mathrm{SD}} \right)+f_{U}}{\overline{\omega}}$ |
| SD | ${f'}_{\mathrm{SD}_{1}}=\frac{f_{\mathrm{SD}_{1}}F_{\mathrm{SD}_{1}}T_{\mathrm{SD}_{1}}{(1+\beta}_{\mathrm{SD}_{1}})}{\bar{\omega}}$  ${m'}_{\mathrm{SD}_{1}}=\frac{f_{\mathrm{SD}_{1}}F_{\mathrm{SD}_{1}}T_{\mathrm{SD}_{1}}(1-\beta_{\mathrm{SD}_{1}})}{\bar{\omega}}$  ${f'}_{\mathrm{SD}_{2}}=\frac{{2f}_{\mathrm{SD}_{2}}F_{\mathrm{SD}_{2}}T_{\mathrm{SD}_{2}}{(1+\beta}_{\mathrm{SD}_{2}})}{\bar{\omega}}$  ${m'}_{\mathrm{SD}_{2}}=\frac{{2f}_{\mathrm{SD}_{2}}F_{\mathrm{SD}_{2}}T_{\mathrm{SD}_{2}}(1-\beta_{\mathrm{SD}_{2}})}{\bar{\omega}}$  ${f'}_{U}=\frac{f_{\mathrm{SD}_{1}}F_{\mathrm{SD}_{1}}\left( 1-T_{\mathrm{SD}_{1}} \right){(1+\beta}_{\mathrm{SD}_{1}})+f_{\mathrm{SD}_{2}}F_{\mathrm{SD}_{2}}\left( 1-T_{\mathrm{SD}_{2}} \right){(1+\beta}_{\mathrm{SD}_{2}})+f_{U}}{\bar{\omega}}$  ${m'}_{U}=\frac{f_{\mathrm{SD}_{1}}F_{\mathrm{SD}_{1}}\left( 1-T_{\mathrm{SD}_{1}} \right)\left( 1-\beta_{\mathrm{SD}_{1}} \right)+f_{\mathrm{SD}_{2}}F_{\mathrm{SD}_{2}}\left( 1-T_{\mathrm{SD}_{2}} \right)(1-\beta_{\mathrm{SD}_{2}})+f_{U}}{\bar{\omega}}$ |
